# Nucleosome sliding by the Chd1 chromatin remodeler relies on the integrity of the DNA duplex

**DOI:** 10.1101/2022.04.19.488687

**Authors:** Sangwoo Park, Taekjip Ha, Gregory D. Bowman

**Affiliations:** Department of Biophysics and Biophysical Chemistry, Johns Hopkins University School of Medicine, Baltimore, MD 21205, USA; Department of Biomedical Engineering, Johns Hopkins University, Baltimore, MD 21205, USA; Howard Hughes Medical Institute, Baltimore, MD 21205, USA; TC Jenkins Department of Biophysics, Johns Hopkins University, Baltimore, MD 21218, USA

## Abstract

Chromatin remodelers use a helicase-type ATPase motor to shift DNA around the histone core. Although not directly reading out the DNA sequence, some chromatin remodelers are biased by DNA sequences, suggesting that they may be sensitive to properties of the DNA duplex. Here, we present a high-throughput method for determining nucleosome positioning in vitro using site-specific DNA cleavage coupled with next-generation sequencing. This method allowed us to systematically test how the introduction of poly(dA:dT) tracts and other perturbations affected the distribution of nucleosomes remodeled by the Chd1 remodeler. We found that Chd1 is sensitive to poly(dA:dT) tracts as short as 3 bp, and that its nucleosome sliding activity is severely perturbed by DNA mismatches and single-nucleotide insertions. These results suggest that remodelers rely on the integrity of duplex DNA for nucleosome sliding. We also discovered that DNA on the nucleosome can shift in the absence of a remodeler when multiple mismatches are placed at superhelix location 2 (SHL2). This DNA movement in response to a disruption of the double helix may explain why SHL2 is the preferred site of engagement by most chromatin remodelers.

## INTRODUCTION

Nucleosomes, the basic packaging unit of eukaryotic chromosomes, limit access to genomic DNA and thus provide a broadly repressive role in transcription and other DNA-related processes (1). Forming a characteristic disk-like shape, nucleosomes consist of ~1.6 superhelical turns of duplex DNA around a histone core, with the DNA backbone tightly coordinated by histones at each inward-facing minor groove (2). Since DNA bending and occlusion antagonizes binding of many DNA-binding factors, nucleosomes must be removed or repositioned to allow DNA to be recognized. Whereas histone-DNA contacts can be disrupted by DNA-tracking motors such as RNA polymerases (3, 4), the regulated repositioning of nucleosomes along DNA in cis requires action of chromatin remodeling enzymes.

Chromatin remodelers can be classified in several structurally and functionally distinct families, yet all use a common Superfamily 2 (SF2)-type ATPase motor to alter nucleosome structure and/or positioning (5). With the exception of INO80, which acts on DNA near the edge of the nucleosome (6–8), most remodelers act at superhelix location 2 (SHL2), an internal site located two helical turns from the central nucleosome dyad (9–13). For remodelers that reposition nucleosomes, the ATPase motor shifts DNA relative to the histone core by distorting DNA at its binding site (14–16). In the first stage of repositioning, the remodeler ATPase pulls one strand of DNA toward the remodeler by one nucleotide (17–19). This strand-specific shift alters the geometry of DNA: the duplex matches an A-form geometry at the ATPase binding site, whereas it shows a distorted base stacking geometry for at least one helical turn outside the binding site (19). The ATPase motor contacts the DNA backbone of both strands, and previous work showed that single-stranded gaps in one DNA strand block nucleosome sliding (8–12). Whether other perturbations of the DNA duplex antagonize nucleosome sliding has not been explored.

Chd1 is a monomeric remodeling factor that repositions (or slides) nucleosomes by acting at SHL2 (12, 19–21). Like several other remodelers (22), the nucleosome sliding activity of Chd1 is sensitive to the sequence of nucleosomal DNA (23–25). In vitro, a commonly used nucleosome positioning sequence is the Widom 601 sequence (26). The Widom 601 sequence possesses an uneven distribution of TpA (TA) dinucleotide steps, with more TA steps on one side of the central dyad (termed the TA-rich side) than the other (the TA-poor side). Chd1 can act at either of the two SHL2 sites on the nucleosome, but has higher activity on the TA-poor side (24, 25). Thus, on 601 nucleosomes flanked by linker DNA on both sides, Chd1 preferentially shifts the histone octamer toward the TA-poor side (23). This bias can be diminshed or reversed by altering the DNA sequence. In particular, inclusion of long (≥16 bp) poly(dA:dT) tracts on the TA-poor side favored nucleosome sliding toward the TA-rich side (23).

Here, we adapted next-generation sequencing (NGS) to determine in vitro positions of nucleosomes after sliding by Chd1. Using the Widom 601 sequence as a base, we determined the sensitivity of nucleosome positioning to poly(dA:dT) tracts of varying lengths and positions. Our results reveal that nucleosome sliding is affected by poly(dA:dT) tracts as short as 3 nt, with the greatest impact when tracts are within one helical turn of the ATPase binding site. We also generated 601-based libraries containing mismatches and single-base insertion. These local disruptions of the DNA duplex had a stronger impact in altering nucleosome sliding activity of Chd1, suggesting that integrity of the doublehelix is important for remodeler-catalyzed translocation of DNA within the nucleosome. We speculate that sensitivity of remodelers like Chd1 to nucleosomal DNA sequence may reflect the stability and energetics of the duplex, providing a means for DNA sequences to bias nucleosome repositioning by remodelers.

## MATERIAL AND METHODS

### Nucleosome library reconstitution

All DNA libraries were custom synthesized (Custom Array Inc.) and amplified by emulsion PCR (27) with unmodified or modified primers. The 601 nucleosome positioning sequence (26) was use as the base for all nucleosomes. To generate the mismatch and insertion/deletion libraries, a pool of single-stranded DNA was first prepared by amplifying with one 5’-phosphorylated primer and then treating with lambda exonuclease, which preferentially digests the phosphorylated strand. Duplex DNA was generated by heat annealing each pool of single-stranded DNA with the complementary strand of canonical single strand 601 DNA, and then purified using miniprep columns (Qiagen). The DNA strands with abasic site were prepared by splint ligation of three DNA fragments purchased from Integrated DNA Technologies (IDT), with one oligo containing the 1,2’-dideoxyribose modification, an abasic site analog. The ligated strands were purified on urea polyacrylamide gels (6%, 19:1 acrylamide and bis, urea 0.5 g/ml) and annealed together to make double strand DNAs in specific top/bottom strand combinations. All nucleosomes were made using *Xenopus laevis* histones that contained H2B (S53C) for site-specific cross-linking with azido phenacyl bromide (APB). Nucleosome libraries were reconstituted by salt-gradient dialysis (28) using a 1:1.2 molar ratio of DNA to histone octamer.

### Nucleosome sliding assays

All nucleosome sliding reaction were done with 150 nM nucleosome, 50 nM Chd1 (for slide-seq experiments) or 50 nM nucleosome, 100 nM Chd1 (for individual sliding experiments) with 2 mM ATP in 1x SlideBuffer (20 mM Tris-HCl pH 7.5, 50 mM KCl, 5 mM MgCl2, 5% sucrose, 0.1mg/ml BSA, and 5 mM DTT). Reactions were stopped at indicated time points with quench buffer (20 mM Tris-HCl pH 7.5, 50 mM KCl, 0.1mg/ml BSA, 5 mM DTT, 5 mM EDTA, and xx μg/μl salmon sperm DNA). Samples were resolved by native PAGE (30%, 60:1 acrylamide and bis) at 4°C.

### Histone-DNA photo-crosslinking and cleavage

After nucleosome sliding by Chd1, nucleosomal DNA was cross-linked to H2B(S53C) labeled with APB (29). Briefly, prior to nucleosome sliding reactions, nucleosomes were labelled with APB in the dark for 2 hours and quenched with xx mM DTT. After sliding, cross-links to DNA were induced by UV illumination for 15 seconds. DNA was processed by incubation at 70°C for 20 min, phenol chloroform extraction, and then ethanol precipitation. Crosslinked DNA was cleaved in 0.1 M NaOH at 90°C for 30 min and quenched with the same volume of 0.1 M HCl. The final fragmented DNA was collected through ethanol precipitation and could be visualized after separation in urea-polyacrylamide gels (8%, 19:1 acrylamide and bis, xx M urea 0.5 g/ml).

### NGS library preparation and sequencing

A schematic of the library preparation and processing is given in Supplementary Figure S1. To preferentially sequence DNAs that were cleaved, a streptavidin pull-down was performed (Invitrogen T1 dynabeads), which removed both biotinylated 5’ end fragments as well as biotinylated uncleaved fragments. After ethanol precipitation of the supernatant, the cleaved single-stranded DNA was filled in to make duplex by Bst 2.0 warmstart polymerase (NEB) at 65°C, and the further cleaned up by AMPure XP beads (Beckman). The final double strand DNA fragments pool was ligated with adapters and amplified with Illumina index primers by using NEBNext Ultra II NGS library prep kit (NEB). The prepared NGS library was sequenced as 150×150 bp pair-end sequencing in the Miseq or Hiseq 2500.

### NGS library data analysis

First, sequencing reads were aligned to the Widom 601 sequence using Bowtie2 software (30). Using the alignment, sequence variations indicated the location of mismatches or insertions/deletions, and a premature truncation in the read indicated the location of cleavage sites. Data were discarded if variation in the 601 sequence or cleavage sites could not be identified. The final sort file, consisting of 601 variant sequences with site-specific cleavage, contained 10-30% of total reads. With the two copies of H2B(S53C), cleavage occurs on both strand, each being 32 nt from the 5’ side of the dyad. Since the sequence variations and cleavage sites must be in the same read, only sequence variations located 3’ to the cleavage site can be detected. We used a linear regression model to combine data from both strands at each position of the 601 to calculate the dyad position. In some cases, such as when the variation was 5’ to a cleavage site, the dyad was determined from one cleavage on one strand. Where noise was significant, the signal was boosted from a simple uniform subtraction of cleavage counts over position. The nucleosome positioning signal was defined as the sum of the cleavage counts at top strand –32 bp and bottom strand +32 bp location.

### NMF score map

First, for each sequence, nucleosome positioning signals was normalized by total reads count and converted into probability measure. Since nucleosome positioning data is nonnegative, we used nonnegative matrix factorization (NMF) method to approximately decompose all positioning data of each library into 3 basis patterns with corresponding weights. For each basis pattern, we mapped associated weight values onto the locations of perturbations at the Widom 601 DNA for each sequence. In the case of multiple scores are assigned onto single locations due to the overlaps of perturbations from different sequences, average score was used.

### Clustering analysis

As the resemblance metric for nucleosome positioning data, the similarity score between two positioning probabilities was defined by exponential of negative Jensen-Shannon divergence of the probability pair. After computing all pair-wise similarity scores, all positioning data of libraries was clustered through Spectral clustering algorithm. The cluster number was determined as the minimal number of getting well isolated clusters through trial errors.

### KL-divergence map

To quantify how much nucleosome positioning signal is deviated from the original Widom 601 case through perturbations, Kullback–Leibler divergence (KL-divergence) metric is used for comparing perturbed positioning probabilities with respect to the original nucleosome positioning. Then the KL-divergence values are mapped on the locations of perturbation at the Widom 601 DNA. In the case of same location have multiple KL-divergence values due to overlaps of perturbation from difference sequences, the averaged value is used.

### Energy landscape analysis

To measure how much DNA perturbations destabilize nucleosomes, we introduced the energy landscape analysis. We converted the nucleosome positioning probability distribution to energy landscape by taking negative natural log. And we computed the how much energy change as introducing DNA perturbations on Widom 601 DNA template. Then these values were mapped onto the DNA template according to the locations and length of the perturbation. For overlapping cases, the average values were taken. At last, the plot was shown in Super Helical Locations (SHL) with respect to 0, +11, +20 bp shifted nucleosomes.

## RESULTS

### Poly(dA:dT) tracts can influence local and global positioning of the Widom 601 sequence

We used next-generation sequencing (NGS) to determine nucleosome positions at high-resolution for a population of different sequences (**Figure 1**). To identify the location of the histone core on DNA, we adapted a site-specific DNA cleavage method described by Bartholomew and coworkers (29). In this method, DNA can be site-specifically cleaved on the nucleosome, 53 bp from the dyad, using a histone core containing H2B(S53C) labelled with the photo-cross-linker azido phenacylbromide (**Figure 1B,C**). After UV cross-linking and processing, each strand of the DNA duplex is cleaved only once on one side of the nucleosome. The sites of cleavage can be determined by NGS, which in turn reveal positioning of the histone core. Since the two H2B(S53C) sites are located 53 bp from the dyad, the locations of these two independent cross-linking sites can be used to determine the position of the nucleosome dyad for each unique sequence in the library (**Figure 1D,E**).

**Figure 1.**
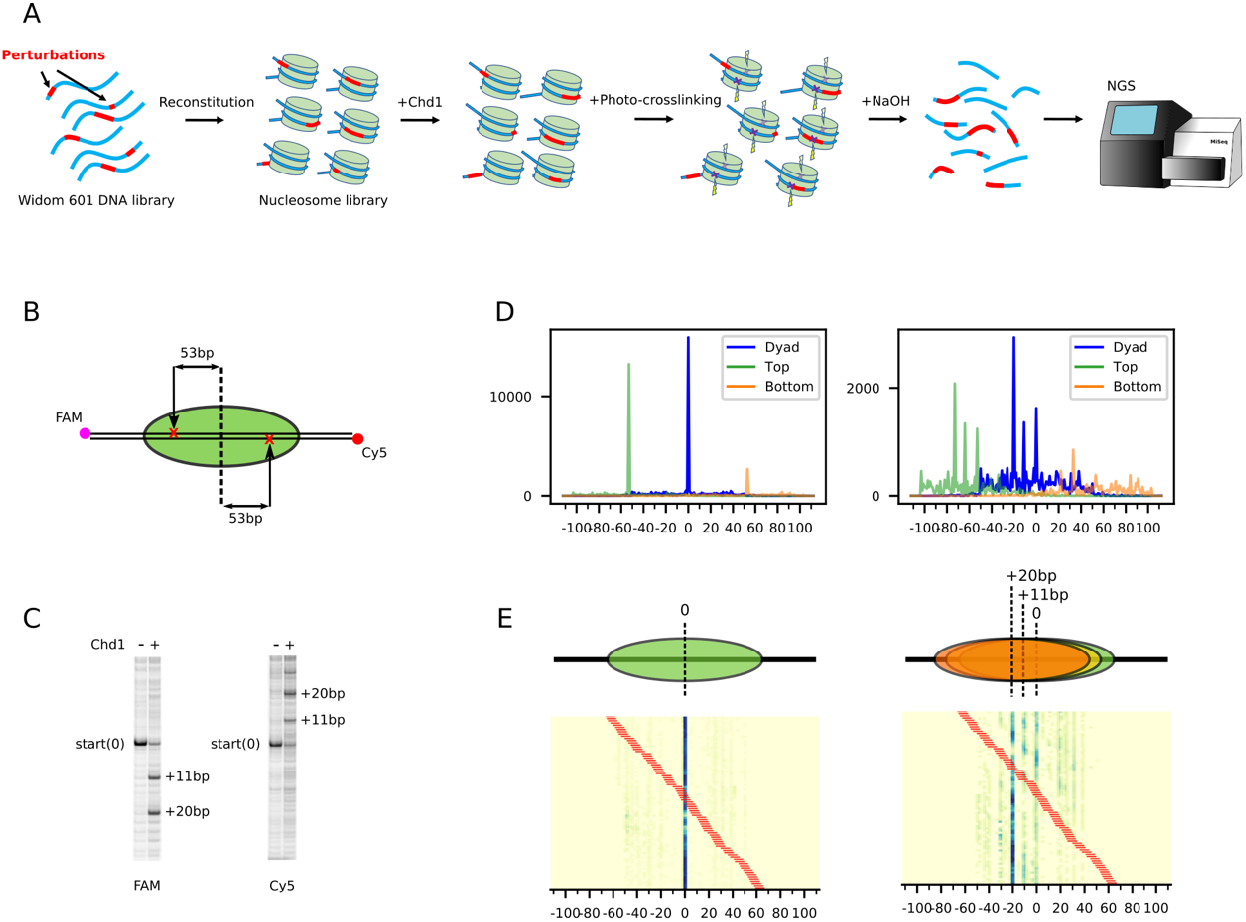
Slide-seq – a strategy using next-generation sequencing to monitor nucleosome sliding. (A) Schematic workflow of slide-seq. After synthesis of a DNA library, based on the Widom 601 positioning sequence, the DNA is incorporated into nucleosomes, and the nucleosomes are then repositioned by the Chd1 remodeler. To determine nucleosome positions, DNA is site-specifically photo-cross-linked to histones, which cleaves the DNA backbone. Nextgeneration sequencing these DNA fragments reveals the locations of DNA cleavages and thus the positions of DNA on the histone core for each distinct sequence. (B) Schematic of the photo-cross-linking sites on the nucleosome. When histone H2B(S53C) is labeled with azido phenacyl bromide, it cross-links to only one strand of the duplex on each side of the nucleosome, 53 bp from the dyad. (C) Example of cleavage products resulting from photo-cross-linking nucleosomes, before and after sliding by Chd1. Here, the photo-crosslinked DNA cleavage was visualized on a urea denaturing gel, scanned separately for FAM (top strand) and Cy5 (bottom strand). (D) Next-generation sequencing data revealing the location of DNA cleavage sides, before (left) and after sliding (right) by Chd1. Green peaks represent cleavage sites on the top strand, orange peaks on the bottom strand, and blue peaks are calculated dyad positions. The DNA sequence for this experiment was the canonical Widom 601 DNA. (E) An example heatmap from the 601 poly(dA:dT) library with 8 bp tract length, before (left) and after sliding (right) by Chd1. Tract location is indicated by the red bars. Cartoons above the heatmaps show most populated nucleosome positions.

To determine the precise positions and lengths where poly(dA:dT) tracts affect activity of Chd1, we generated a library where every position of the canonical 145 bp 601 sequence is the starting point for a tract of 3 to 15 bp. After amplifying the library using emulsion PCR (27), nucleosomes were assembled using the standard salt gradient dialysis method, then UV cross-linked and processed for sequencing, either before or after repositioning by yeast Chd1. We found that a number of poly(dA:dT) tracts strongly affected nucleosome positioning during salt gradient dialysis, prior to repositioning by Chd1. We first describe how the lengths and locations of poly(dA:dT) tracts altered the expected sharp dyad positioning of the canonical Widom sequence. We then focus on how the length and position of poly(dA:dT) tracts influenced nucleosome repositioning by Chd1.

After generating a library of nucleosomes containing poly(dA:dT) tracts, dyad positions were mapped and sorted based on poly(dA:dT) position and length (**Supplementary Figures S1 and S2, Supplementary Table 1**). Clustering analysis Based on the similarity of nucleosome positions and perturbation patterns, the poly(dA:dT) library data was clustered into 6 groups (**Supplementary Figure S3**). To visualize where and how poly(dA:dT) tracts most strongly affected nucleosome positioning, we used a machine learning analysis method, called None-negative matrix factorization (NMF) (31) (**Figure 2**). With this method, the complex pattern of dyad positions for each sequence could be linearly decomposed into a few basis patterns with corresponding weights. Here, these weights are referred to as the NMF scores for each pattern. For a given length of poly(dA:dT) tract (3-mer, 4-mer, etc), all NMF scores were mapped onto the Widom 601 sequence according to tract location. This mapping construction produced a ‘geographic’ heat map indicating where poly(dA:dT) tracts of different lengths were most prevalent for each dyad pattern. We found three predominant dyad patterns, which we refer to as “clean”, “noisy”, and “split” (**Figure 2A**). For each pattern, high NMF scores indicate the locations and lengths of poly(dA:dT) tracts that produced a similar distribution.

**Figure 2.**
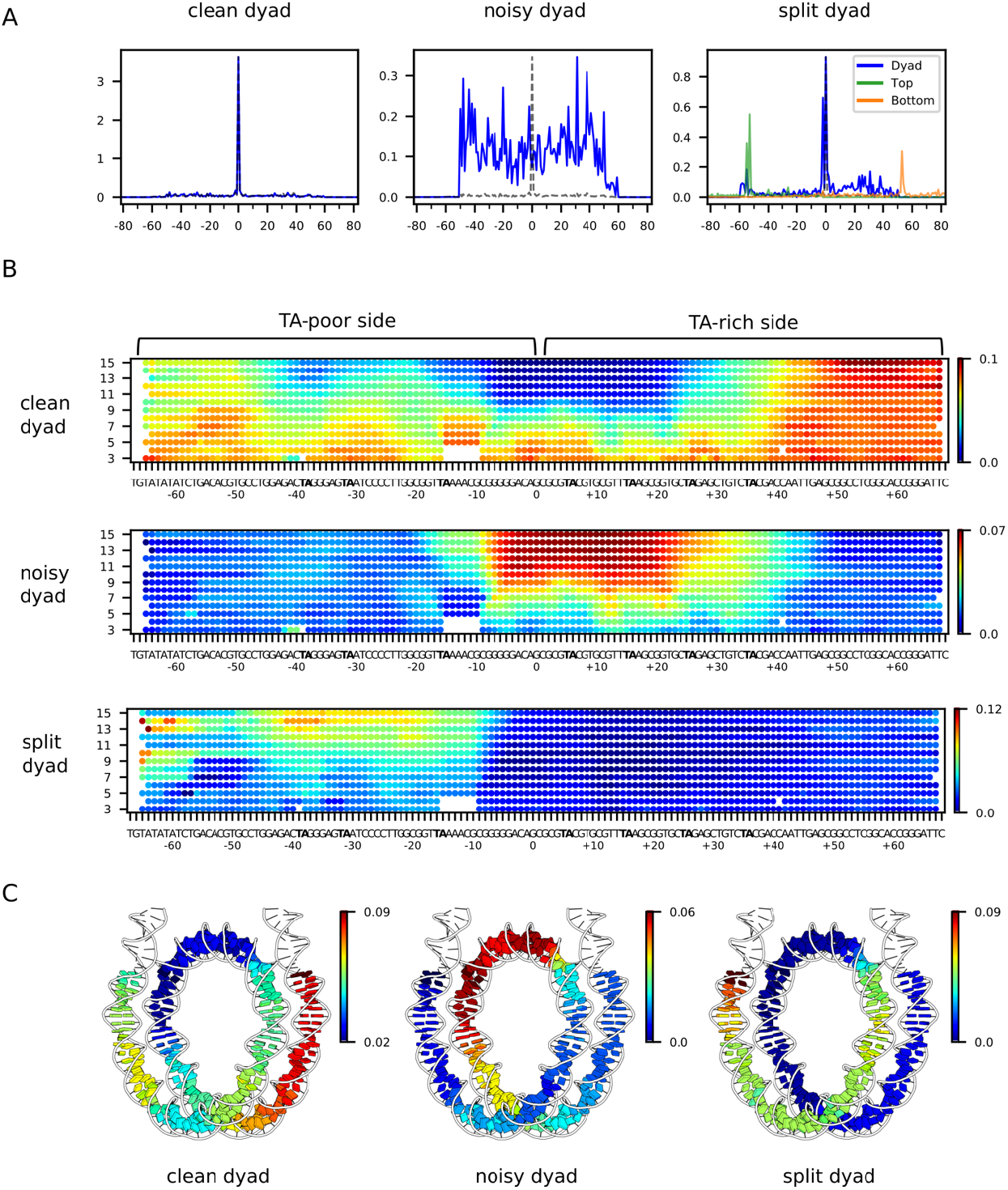
NMF analysis for nucleosome positioning on the 601 poly(dA:dT) library before sliding reveals the asymmetric nature of nucleosome stability on Widom 601 library. (A) For nonnegative matrix factorization (NMF) analysis, all nucleosome positioning signals of the 601 poly(dA:dT) library were categorized based on three basis patterns: (1) a single dominant peak as expected for the 601 (clean dyad), (2) a noisy pattern (noisy dyad), and (3) doublet peaks for the top strand and calculated dyad (split dyad). Calculated dyad positions are shown in blue, and H2B(S53C) cleavage sites in orange and green. Y-axis values are relative frequencies. (B) A geographic heat map of NMF scores for each dyad pattern. Poly(dA:dT) tract lengths are indicated on the left. (C) NMF scores for nucleosomes containing 10 bp poly(dA:dT) tracts mapped onto the nucleosome structure (6WZ5).

The clean dyad pattern resembles the canonical Widom 601, and therefore shows where poly(dA:dT) tracts did not disrupt the 601-directed positioning of the histone core. This pattern was enriched in the shorter tracts, and those located 40 bp or more away from the dyad on the TA-rich side. The noisy dyad pattern, in contrast, had a broad distribution of the dyad compared with the unique 601-directed dyad position (**Figure 2B**), which indicates that these poly(dA:dT) tracts interfered with positioning by the rest of the 601 sequence. The noisy pattern was enriched in the longer poly(dA:dT) tracts (>8 bp) and extended over a ~40 bp segment that included the dyad and was mostly on the TA-rich side of the 601 sequence. The TA steps have been shown to be important sequence elements that allow nucleosomal DNA to wrap more favorably (32), and therefore the elimination of these sequence elements by overlapping poly(dA:dT) tracts may explain disruptions of the canonical 601 dyad position. However, the noisy dyad pattern also had strong NMF scores with some shorter, specifically placed poly(dA:dT). In particular, poly(dA:dT) tracts as short as 6 bp were enriched at on the TA-rich side on SHL1 and SHL2, between positions [8:13] and [19:24]. Interestingly, several of these sites correspond with where the minor groove of DNA would be outward-facing and therefore widest on canonical 601 nucleosomes, suggesting that these poly(dA:dT) tracts may interfere with positioning due to their intrinsically narrow minor groove width (33).

The split dyad pattern arose from a 2 nt shift in H2B cross-link in only the top strand, which is on the TA-poor side of the 601 sequence. This pattern corresponds to a previously observed shift of entry-side DNA upon binding of Chd1 to the TA-poor side of Widom 601 nucleosomes (14). As shown by cryo-EM work, entry-side DNA is shifted toward the dyad upon binding of remodeler ATPases to SHL2 in a nucleotide-free or ADP-bound state (17–19). Here, a similar shift appears to be stimulated with poly(dA:dT) sequences located on the TA-poor side of the Widom 601 sequence (**Figure 2B,C**). This pattern suggests that poly(dA:dT) tracts on the TA-poor side do not affect positioning of the dyad per se, but instead cause a slight shift of entry-side DNA, without the need for binding of a remodeler ATPase at SHL2.

### Poly(dA:dT) tracts antagonize nucleosome sliding immediately downstream of the Chd1 binding site

The poly(dA:dT) tract library also showed where different lengths and positions of tracts affected nucleosome positioning by the Chd1 remodeler (**Figure 3A, Supplementary Figure S4**). Previously, Chd1 was shown to preferentially shift 40N40 nucleosomes made with the Widom 601 sequence toward the TA-poor side (23). After nucleosome sliding by Chd1, we obtained a similar distribution for many nucleosomes in the poly(dA:dT) library, with the strongest peak 20 bp away from the starting position. However, many lengths and positions of poly(dA:dT) tracts influenced nucleosome positioning by Chd1.

**Figure 3.**
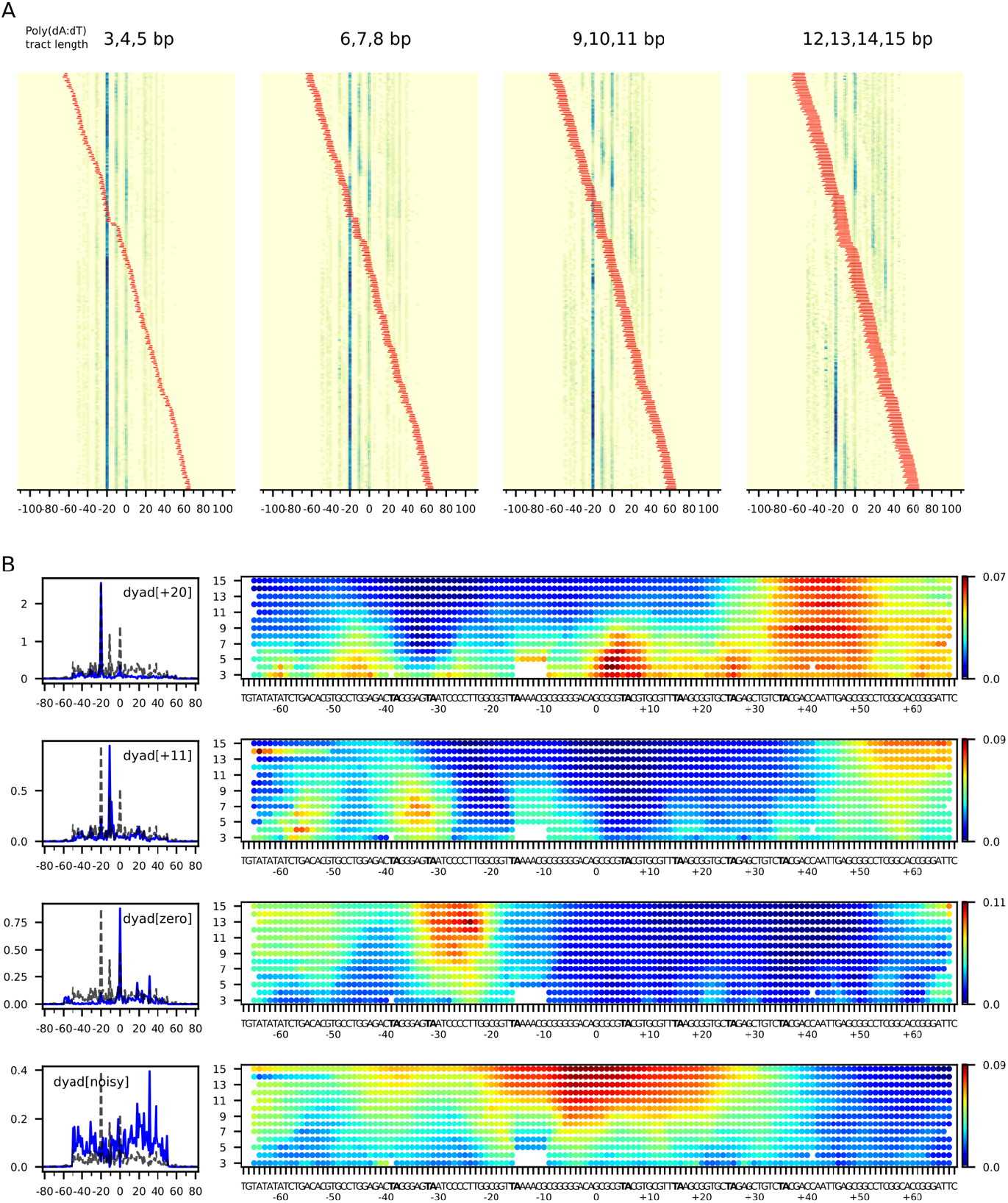
Poly(dA:dT) tracts alter the distribution of nucleosomes after sliding by Chd1. (A) Heatmaps for estimated nucleosome dyads signals after sliding by Chd1. The sequences are grouped according to the poly(dA:dT) tract lengths, 3-5 bp, 6-8 bp, 9-11 bp and 12-15 bp, and sorted by tract locations (red). (B) Dyad patterns (left) and NMF heatmaps (right) for the four dominant classes of calculated dyads. In the dyad patterns, the dotted lines indicate the dyad distribution for Chd1 sliding of the canonical Widom 601 nucleosome.

We generated NMF plots based on four common nucleosome repositioning patterns: (1) a 20 bp shift toward the TA-poor side (dyad[+20]), similar to the canonical Widom 601; (2) a predominant 11 bp shift toward the TA-poor side (dyad[+11]); (3) no shift (dyad[zero]); and (4) a noisy dyad pattern (**Figure 3B**). The noisy dyad pattern correlated with the positions and lengths of polyA(dA:dT) tracts that disrupted the canonical 601 positioning prior to remodeling by Chd1 (noisy dyad, **Figure 2**).

Since Chd1 is known to bind and translocate DNA when bound to SHL2, we can interpret the influence of the poly(dA:dT) tracts by their position relative to the remodeler binding site. Poly(dA:dT) tracts that favored the dyad[+20] pattern, and therefore had little effect on Chd1 activity, were primarily located on the opposite side of the nucleosome. In contrast, the sequences that interfered with the expected Chd1 sliding pattern were closer to the SHL2 site on the TA-poor side.

Interestingly, the poly(dA:dT) sequence that appeared to block sliding by Chd1 (dyad[zero]) were distributed asymmetrically with respect to the ATPase motor binding site. Like other remodelers, the Chd1 ATPase motor acts at SHL2, with direct contacts to both strands of DNA between 16-24 bp from the dyad (19, 34, 35). Poly(dA:dT) tracts that increased the dyad[zero] population primarily occupied regions on the entry side (as opposed to the dyad side) of the ATPase binding site. Since Chd1 shifts DNA at SHL2 toward the dyad (12), the entry side corresponds with downstream DNA that will be pulled onto the ATPase binding site. Several poly(dA:dT) tracts that increased the dyad[zero] species were located completely outside the binding site (e.g. A_8_[-36:-29], A_8_[-35:-28], and A_8_[-34:-27]; **Supplementary Table 2**). This indicates that the action or efficiency of the remodeler ATPase motor was most strongly affected by the sequence of DNA that was immediately downstream of the binding site, poised to be shifted toward the remodeler at SHL2.

The poly(dA:dT) tracts that favored the dyad[+11] pattern were predominantly located at SHL3. These poly(dA:dT) tracts allowed for an initial 11 bp shift of the histone core, but then reduced or blocked further movement. To visualize how these tracts may affect the remodeler at SHL2, we mapped the NMF heatmaps onto the nucleosome before and after repositioning (**Figure 4**). After an 11 bp shift, the tracts enriched in the dyad[+11] pattern were also asymmetrically positioned on the remodeler binding site, with a bias toward the entry side. These experiments therefore indicate that poly(dA:dT) tracts primarily interfered with nucleosome sliding when they overlapped with the ATPase binding site and were within one helical turn of the binding site on the downstream side.

**Figure 4.**
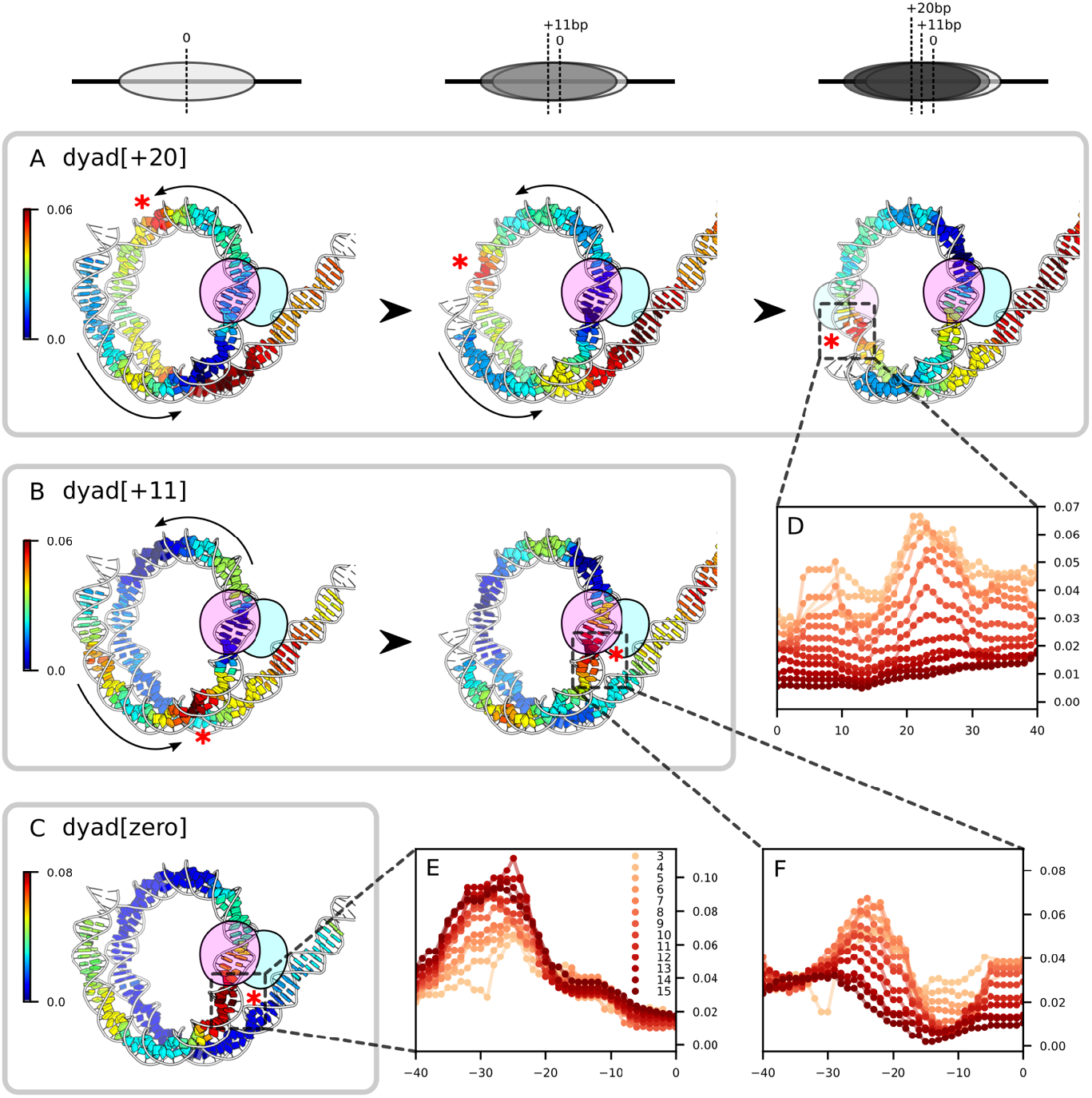
Poly(dA:dT) tracts on the downstream side of Chd1 binding sites strongly interfere Chd1 sliding. (A) NMF scores for the 8 bp poly(dA:dT) tracts with the dyad[+20] pattern, mapped onto a nucleosome-Chd1 complex solved by cryo-EM (7TN2). Shown are NMF scores at three translational positions of the 601 sequence: at the starting position for 601 (right), after a +11 bp shift (middle), and after a +20 bp shift (left). (B) NMF scores for the 8 bp poly(dA:dT) tracts with the dyad[+11] pattern, mapped onto a nucleosome-Chd1 complex (7TN2). Shown are NMF scores at two translational positions of the 601 sequence: at the starting position for 601 (right), and after a +11 bp shift (left). (C) NMF scores for the 8 bp poly(dA:dT) tracts with the dyad[zero] pattern, mapped onto a nucleosome-Chd1 complex (7TN2). Shown are NMF scores at the starting position for 601. (D) NMF scores for all tract lengths with the dyad[+20] pattern, corresponding to the region around the opposing SHL2 after a +20 bp shift. (E) NMF scores for all tract lengths with the dyad[zero] pattern, corresponding to the region around SHL2. (F) NMF scores for all tract lengths with the dyad[+11] pattern, corresponding to the region around SHL2 after a + 11 bp shift.

### Nucleosome sliding by Chd1 is sensitive to the integrity of downstream DNA

Given that longer poly[dA:dT] tracts showed a stronger interference with sliding activity, we wondered whether the remodeler was sensitive to stability of the DNA duplex. We therefore generated two new libraries designed to site-specifically alter DNA duplex structure: (1) a mismatched library, where non-A•T bp were replaced with an A on the top strand or T on the bottom strand, yielding six different types of mismatches; and (2) an insertion library, where an additional A nucleotide was introduced on either the top or the bottom strand along the Widom 601 sequence. Both of these disruptions relied upon having one strand maintain the canonical 601 sequence, and the other strand possess a discrete difference. To achieve this, the canonical and altered 601 sequences were separately amplified with PCR, and then one of the two strands selectively destroyed with lambda exo, which targets 5’ phosphorylated substrates (36). Annealing the remaining strands together (e.g. the top strand from amplification of the canonical 601 sequence and bottom strand of amplification of a variant 601 sequence) produced a duplex that properly base-paired except at the site of the sequence difference (**Supplementary Figure S5**). With these modified duplexes, nucleosomes were generated through salt gradient dialysis and then cross-linked and sequenced before and after sliding by Chd1.

Before sliding by Chd1, the mismatch library of 601 nucleosomes showed similar trends as those of the poly(dA:dT) library. Whereas a majority of mismatch positions yielded a sharp dyad signal like the canonical 601 (‘clean dyad’), sites on and around the dyad gave a broader distribution of dyad positions (‘noisy dyad’, **Supplementary Figures S6-S8, Supplementary Table 3**). Unlike the poly(dA:dT) library, though, the most disruptive sites were on the TA-poor side of the dyad. Interestingly, the mismatches also produced a ‘split dyad’ signal, which, like the poly(dA:dT) library, was due to a cross-linking doublet on the TA-poor side of the 601 sequence. For the poly(dA:dT) tracts, the cross-linking doublet was most prominent with long tracts spanning SHL2 to SHL4. For the mismatches, however, the cross-linking doublet was limited to those at SHL2, located 16-25 bp from the dyad (**Supplementary Figure S8**). Interestingly, this is precisely where chromatin remodelers distort nucleosomal DNA to allow absorption of an extra nucleotide, which stimulates movement of entry DNA toward the remodeler (14, 17, 19).

The mismatch library was then remodeled with Chd1, and the dyad positions calculated as before. For the mismatch library, a similar trend was observed as for the poly[dA:dT] library: some nucleosomes were shifted 20 bp toward the TA-poor side (dyad[+20]), some were predominantly shifted only 11 bp (dyad[+11]), and others remained at the starting position (dyad[zero]) (**Figure 5, Supplementary Figure S9**). Strong effects were observed for two or more consecutive mismatches, with single mismatches having only modest impacts. The ≥2 bp mismatches located on the TA-poor side, 32-37 bp from the dyad, gave an increase in the dyad[+11] species, whereas those 18-29 bp from the dyad prevented nucleosomes from shifting from the starting dyad[zero] position. As seen for the poly[dA:dT] tracts, mismatches just outside the ATPase binding site strongly interfered with sliding (e.g. M2[-29:-28], **Supplementary Table 4**), indicating that the integrity of the downstream DNA duplex was important for sliding.

**Figure 5.**
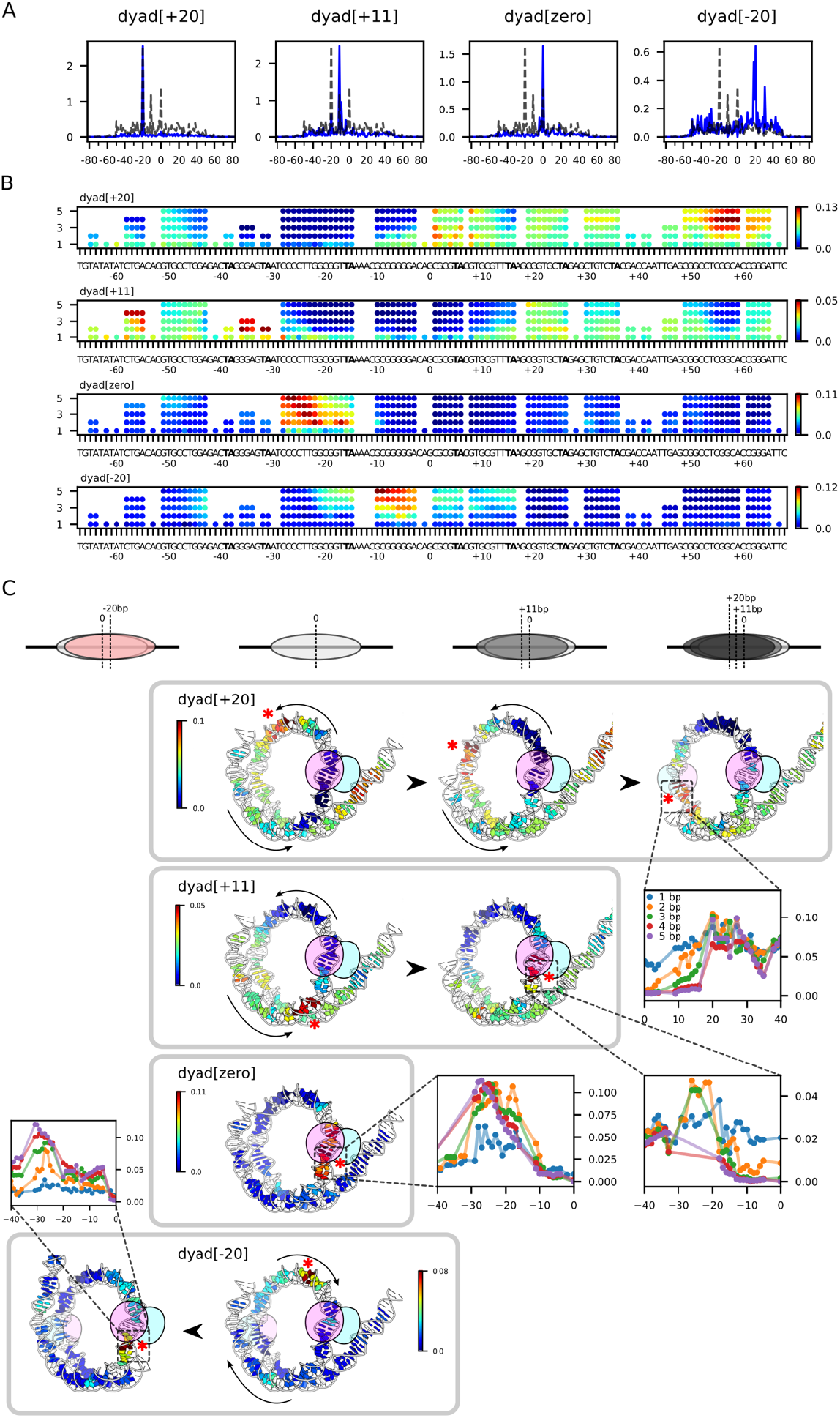
Mismatches bias the distribution of nucleosome positions after sliding by Chd1. (A) Four basis dyad patterns observed after sliding by Chd1, which were used for NMF scoring (blue). The distribution of the canonical 601 is shown with dotted lines. (B) NMF scores mapped onto the Widom 601 DNA sequence based on the dyad[+20], dyad[+11], dyad[zero], and dyad[-20] patterns. (C) NMF scores for 3 bp mismatches mapped onto a Chd1-nucleosome complex (7TN2) at different translational positions, as shown in Figure 4.

The mismatch library also produced a group of nucleosomes where Chd1 shifted the dyad toward the TA-rich side of the 601 sequence (dyad[-20]; **Figure 5**). The shift toward the TA-rich side corresponded with mismatches between the dyad and +10 on the TA-poor side. These TA-rich species are consistent with the notion that Chd1 dynamically acts on both sides of the nucleosome, with the distribution of nucleosome positions revealing where the remodeler was most efficient/active. Once the histone core is shifted toward the TA-rich side, the mismatches that were previously between zero and +10 become shifted onto SHL2 and SHL3 of the TA-poor side, thereby blocking remodeler action on that side and thus enriching nucleosomes on the TA-rich side of the 601 sequence. As seen from the heatmaps, a similar behavior of shifting toward the TA-rich side was also apparent in the poly(dA:dT) library (**Figure 3A**). However, the nucleosome dyad patterns for the poly(dA:dT) library were not as strongly affects as the mismatch library, perhaps signifying the stronger impact of the mismatches disrupting the integrity of the DNA duplex.

A similar trend in nucleosome positioning was observed with the 1-nt insertion library (**Figure 6, Supplementary Figure S10, Supplementary Tables 5&6**). Before sliding by Chd1, a split dyad signal was observed with an insertion between 13 and 14 nt from the dyad (I_1_[13^14]; **Supplementary Figure S11, Supplementary Table 5**). Like the mismatch library, this shifted peak was only observed on one side of the nucleosome, and therefore appears to represent a shift in DNA at the cross-linking site rather than the dyad. Unlike the mismatch and poly(dA:dT) libraries, however, this shift was on the TA-rich side of the nucleosome and effectively ratcheted entry/exit DNA at the cross-linking site away from, rather than toward, the dyad. Although it is not presently clear why an insertion at this site but not others stimulates a shift in cross-linking, such a shift could be explained by a loss of 1 bp at the TA-rich SHL2 site. In nucleosome crystal structures, SHL2 has been shown to accommodate 10 or 11 bp, with 11 bp preferred by the 601 sequence (16, 37, 38). Therefore, if the additional unpaired base at this location favored 10 bp at SHL2 instead of 11 bp, it could explain the 1 nt shift at the cross-linking site, ~40 bp away.

**Figure 6.**
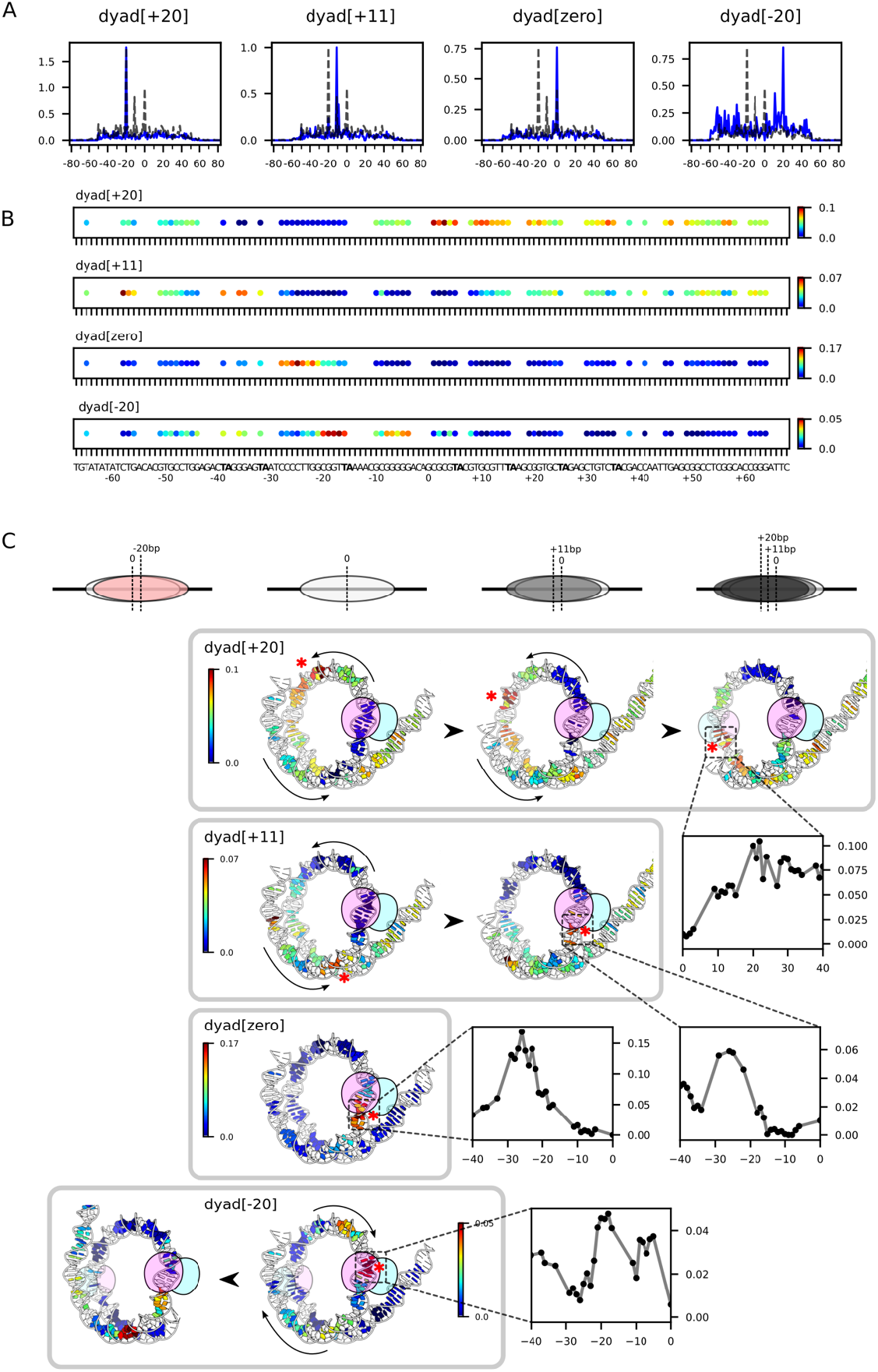
Single-nucleotide insertions alter the distribution of nucleosomes repositioned by Chd1. (A) Four basis dyad patterns observed after sliding by Chd1, which were used for NMF scoring (blue). The distribution of the canonical 601 is shown with dotted lines. (B) NMF scores mapped onto the Widom 601 DNA sequence based on the dyad[+20], dyad[+11], dyad[zero], and dyad[-20] patterns. (C) NMF scores mapped onto a Chd1-nucleosome complex (7TN2) at different translational positions, as shown in Figure 4.

After sliding by Chd1, single nucleotide insertions located between +40 and +33 bp from the dyad blocked the dyad[+20] species but enriched for dyad[+11], whereas those located +29 to +19 blocked both the dyad[+20] and dyad[+11]. Although an inserted base would be expected to interfere with direct binding of the ATPase motor to its target site, the interference from 1-nt insertions outside the binding site (e.g. 28_29 from the dyad) supports the idea that disruption of DNA duplex integrity immediately downstream of the binding site antagonizes nucleosome repositioning by Chd1.

### The importance of base-stacking is different for the two strands of nucleosomal DNA

The results above suggest that weakening or disruption of DNA duplex integrity antagonizes nucleosome sliding by Chd1. Based on these data, we wondered whether base stacking was equally important in both DNA strands of the duplex. To address this question, nucleosomes were created that contained abasic (AP) sites. Unlike previous work using single-stranded DNA gaps (10–12), here the sugar-phosphate backbone of both strands remained intact, with only removal of a base on one strand. Abasic sites were targeted to the middle of the ATPase binding site, 20 bp from the dyad. At the ATPase binding site, the DNA strand oriented 5’-3’ toward the dyad is called the tracking strand, whereas the other strand is called the guide strand (16). Nucleosomes were generated with an abasic site on the tracking strand only, the guide strand only, or simultaneously on both strands. Unlike the 40N40 nucleosomes used for sequencing libraries, these nucleosomes were end-positioned (0N80), with flanking DNA only on one side. As previously described (39), Chd1 requires entry-side DNA for sliding activity, and therefore only acts at the SHL2 proximal to the 80-bp linker DNA. Nucleosome repositioning was monitored by native gels, where nucleosomes migrate more slowly as they move from end- to more centrally-positioned sites.

Compared to the control, nucleosomes containing abasic sites were shifted significantly more slowly by Chd1 (**Figure 7**). While both abasic sites perturbed Chd1 activity, the abasic site on the tracking strand was significantly more deleterious (~60-fold) than the guide strand (~20-fold). Having both abasic sites together essentially abolished sliding. These impacts of removing a single base at the remodeler binding site suggests that base stacking is indeed important for remodeler action. Further, the asymmetry in activity toward abasic sites on the two strands suggests that stacking plays a more important role in the tracking strand. Like other remodelers, Chd1 binding in the nucleotide-free state can shift the tracking strand by 1 nt while keeping the guide strand in place at its binding site (19). This tracking-only shift alters duplex geometry that is likely important for translocation of DNA toward the dyad. The sensitivity to an abasic site on the tracking correspond with this first critical step in nucleosome repositioning.

**Figure 7.**
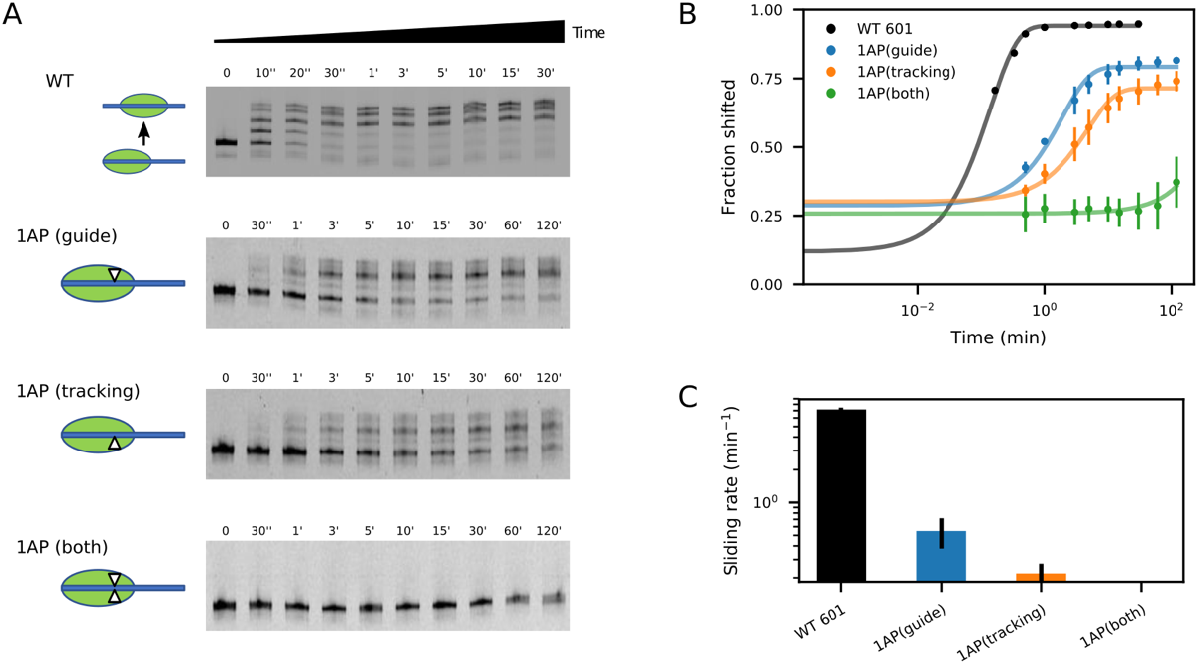
Abasic sites at SHL2 interfere with sliding by Chd1. (A) Nucleosome sliding experiments carried out with 100 nM Chd1, 50 nM FAM-labeled 80N0 nucleosomes, and analyzed by native acrylamide gels. Canonical 601 nucleosomes are shown at the top, and the bottom three gels show nucleosomes with abasic sites at SHL2 on either the guide strand, tracking strand, or both strands (triangles). These gels are representative of three replicates. (B) Quantification of nucleosome sliding gels, as shown in panel A. Shown are the means plus standard deviations from three replicates. Lines show the best single exponential fits. (C) Nucleosome sliding rates: canonical 601 = 7.16 min^-1^; 1 AP[guide] = 0.55 min^-1^; 1 AP[tracking] = 0.22 min^-1^; and 1 AP[both] ≤ 0.01 min^-1^. Shown are the mean rates with standard deviations, calculated from fits for individual experiments.

## DISCUSSION

In this work, we show that localized destabilization of duplex DNA antagonizes nucleosome sliding by the Chd1 remodeler. Biochemically, we show that nucleosome sliding is perturbed when base stacking is perturbed at the ATPase binding site (**Figure 7**). Interestingly, however, with NGS we found that DNA disruptions that affected nucleosome redistributions were enriched around SHL3, outside of the ATPase binding site. We believe that DNA disruptions outside the binding site could affect nucleosome repositioning in two ways. First, since remodelers can act on either side of the nucleosome, DNA disruptions on one side will increase the likelihood of nucleosome sliding on the other side. Due to the two-fold nature of the nucleosome, action of a remodeler on the two sides shifts DNA in opposite directions. Therefore, when one SHL2 site has a block, action at the other SHL2 site will shift that block from SHL2 to SHL3 (**Supplementary Figure S12**). We therefore expect that DNA disruptions that block sliding only at SHL2 could redistribute back toward SHL3 due to sliding on the other side of the nucleosome.

Another possibility, however, is that a DNA disruption at SHL3 directly interferes with nucleosome sliding by Chd1 (**Figure 8**). In support of this idea, a recent cryo-EM structure revealed that the DNA distortion created by Chd1 in the nucleotide-free state alters the geometry of downstream DNA (19). The altered geometry suggests that downstream DNA adopts higher energy conformations in response to distortions at SHL2. Since downstream DNA would be destabilized by mismatches and single-base insertions, we suspect that particular constrained structures of DNA created by the remodeler ATPase could be instrumental in shifting the DNA duplex past the histone core.

**Figure 8.**
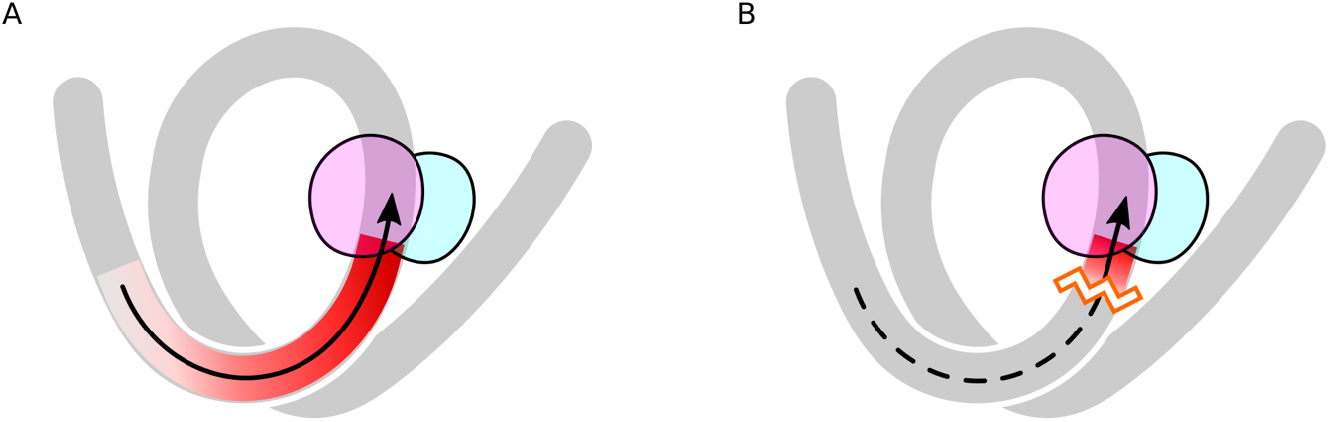
Distortions of downstream DNA impair Chd1 chromatin remodeling. (A) During nucleosome sliding, the ATPase motor of a chromatin remodeler distorts DNA, which is initiated at its SHL2 binding site and extends toward the entry side (red). This distortion is believed to be coupled to translocation of entry DNA toward the remodeler. (B) Destabilization of the DNA duplex just outside the ATPase binding site may block propagation of stress or tension initiated by the remodeler at SHL2, which in turn could disrupt the sliding reaction.

Similarly to mismatches and single-base insertions, interference with nucleosome sliding was also observed when DNA downstream of the remodeler binding site contained poly(dA:dT) tracts. Since they favor a specific structure with narrower minor grooves, poly(dA:dT) tracts may disfavor DNA conformations induced by Chd1, which specifically widens the minor groove of downstream DNA in the nucleotide-free state (19). Poly(dA:dT) tracts are readily accommodated in nucleosomes (40, 41) and their lengths and frequencies throughout eukaryotic genomes vary in a species-specific manner (42, 43). Consistent with previous work (23), site-specific incorporation of poly(dA:dT) tracts near SHL2 on the TA-poor side of centered 601 nucleosomes biased the remodeled distribution toward the TA-rich side. As shown here, the strength of poly(dA:dT) interference is coupled to tract location and length, suggesting a sequence-based means for tuning nucleosome sliding activity and biasing patterns of nucleosome arrays. As shown in a prior study, DNA sequence can influence the distribution of remodeled nucleosomes in vitro (22). We speculate that, for some classes of remodelers, the sequence dependence may localize to the segment of DNA distorted by the ATPase motor, where constraints from both the remodeler and histone core may amplify sequence-dependent properties of the DNA duplex.

An unexpected finding was that, in the absence of a remodeler, DNA mismatches at SHL2 gave rise to a “split dyad” pattern, which corresponded to a DNA shift at one SHL5 cross-linking site, ~30 nt away. Interestingly, this DNA shift is similar to that stimulated by remodeler binding in the nucleotide-free and ADP-bound forms (14). By altering the DNA structure at SHL2, remodelers initially stimulate a concerted corkscrew shift of entry-side DNA by ~1 bp toward SHL2 (17–19). Here, a similar shift of entry DNA by mismatches alone suggests that SHL2 may naturally accommodate additional DNA, which would enable a corresponding corkscrew shift of the DNA duplex. SHL2 and SHL5 are known for allowing for a 1 bp variation in duplex length (16, 37, 38), though the shift seen here with mismatches would necessitate absorption of 1 bp more than nucleosomes with unperturbed DNA. These findings suggest that SHL2 intrinsically couples local DNA distortions with a corkscrew shift of the adjacent DNA segment, which may explain why SHL2 is the preferred site of engagment for many chromatin remodelers.

## ACCESSION NUMBERS

Sequencing data has been deposited in the GEO database with accession number GSE198440 (private token: yfcrkwiojdgrzmn).

## SUPPLEMENTARY DATA

Supplementary Data are available at NAR online.

## ACKNOWLEDGEMENT

We thank Ilana Nodelman and Jessica Winger for helpful discussions.

## SUPPLEMENTARY TABLE LEGENDS

Supplementary Table 1. Sequence reads for top-strand cross-links (green), bottom-strand crosslinks (orange), and predicted nucleosome dyads (blue) for nucleosomes in the poly(dA:dT) library, prior to nucleosome sliding by Chd1.

Supplementary Table 2. Sequence reads for top-strand cross-links (green), bottom-strand crosslinks (orange), and predicted nucleosome dyads (blue) for nucleosomes in the poly(dA:dT) library, after nucleosome sliding by Chd1.

Supplementary Table 3. Sequence reads for top-strand cross-links (green), bottom-strand crosslinks (orange), and predicted nucleosome dyads (blue) for nucleosomes in the mismatch library, prior to nucleosome sliding by Chd1.

Supplementary Table 4. Sequence reads for top-strand cross-links (green), bottom-strand crosslinks (orange), and predicted nucleosome dyads (blue) for nucleosomes in the mismatch library, after nucleosome sliding by Chd1.

Supplementary Table 5. Sequence reads for top-strand cross-links (green), bottom-strand crosslinks (orange), and predicted nucleosome dyads (blue) for nucleosomes in the insertion library, prior to nucleosome sliding by Chd1.

Supplementary Table 6. Sequence reads for top-strand cross-links (green), bottom-strand crosslinks (orange), and predicted nucleosome dyads (blue) for nucleosomes in the insertion library, after nucleosome sliding by Chd1.

