## SupplementaryFigures for "Nucleosome sliding by the Chd1 chromatin remodeler relies on the integrity of the DNA duplex"

### SUPPLEMENTARY FIGURES

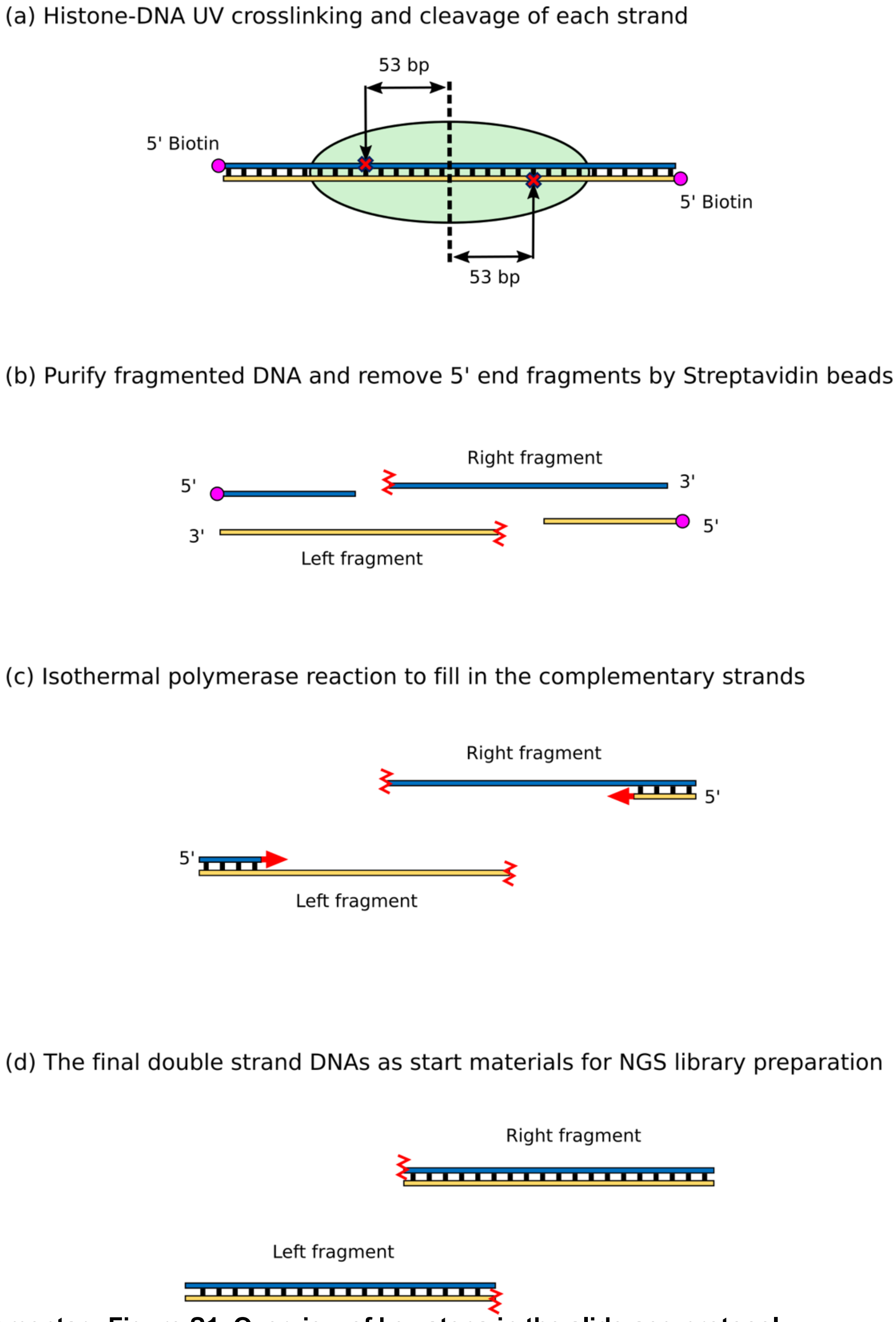


**Supplementary Figure S1. Overview of key steps in the slide-seq protocol.**

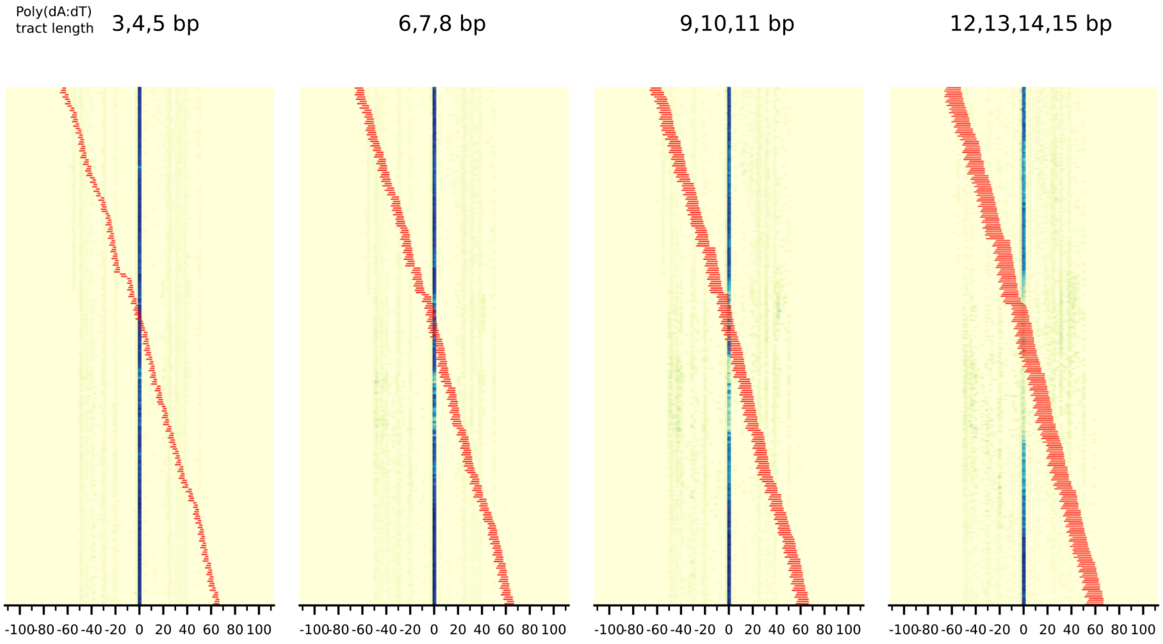


**Supplementary Figure S2. Heatmaps for estimated nucleosome positioning signals on poly(dA:dT) library before sliding.**

The sequences were grouped according to the lengths and locations of poly(dA:dT) tracts (red).

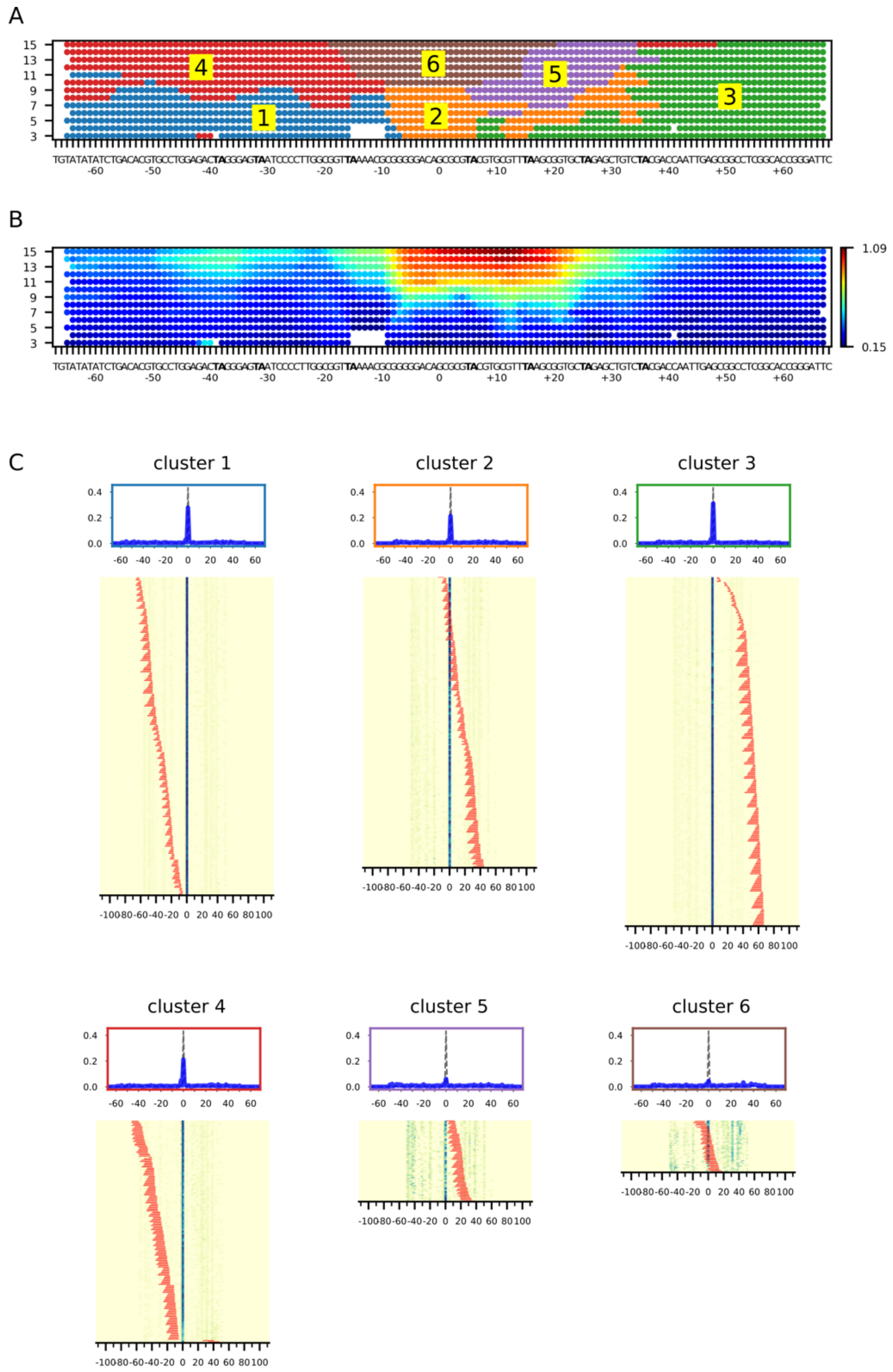


**Supplementary Figure S3. Clustering analysis of the nucleosome positioning on poly(dA:dT) library before sliding by Chd1.**

(A) Based on the similarity of nucleosome positions and perturbation locations, poly(dA:dT) library data was clustered into 6 groups. (B) A KL-divergence heatmap shows the most sensitive area in the Widom 601 DNA by perturbations. (C) Dyad positions for each poly(dA:dT) tract are shown as heatmaps according to each cluster. Red bars indicate the length and position of each tract.

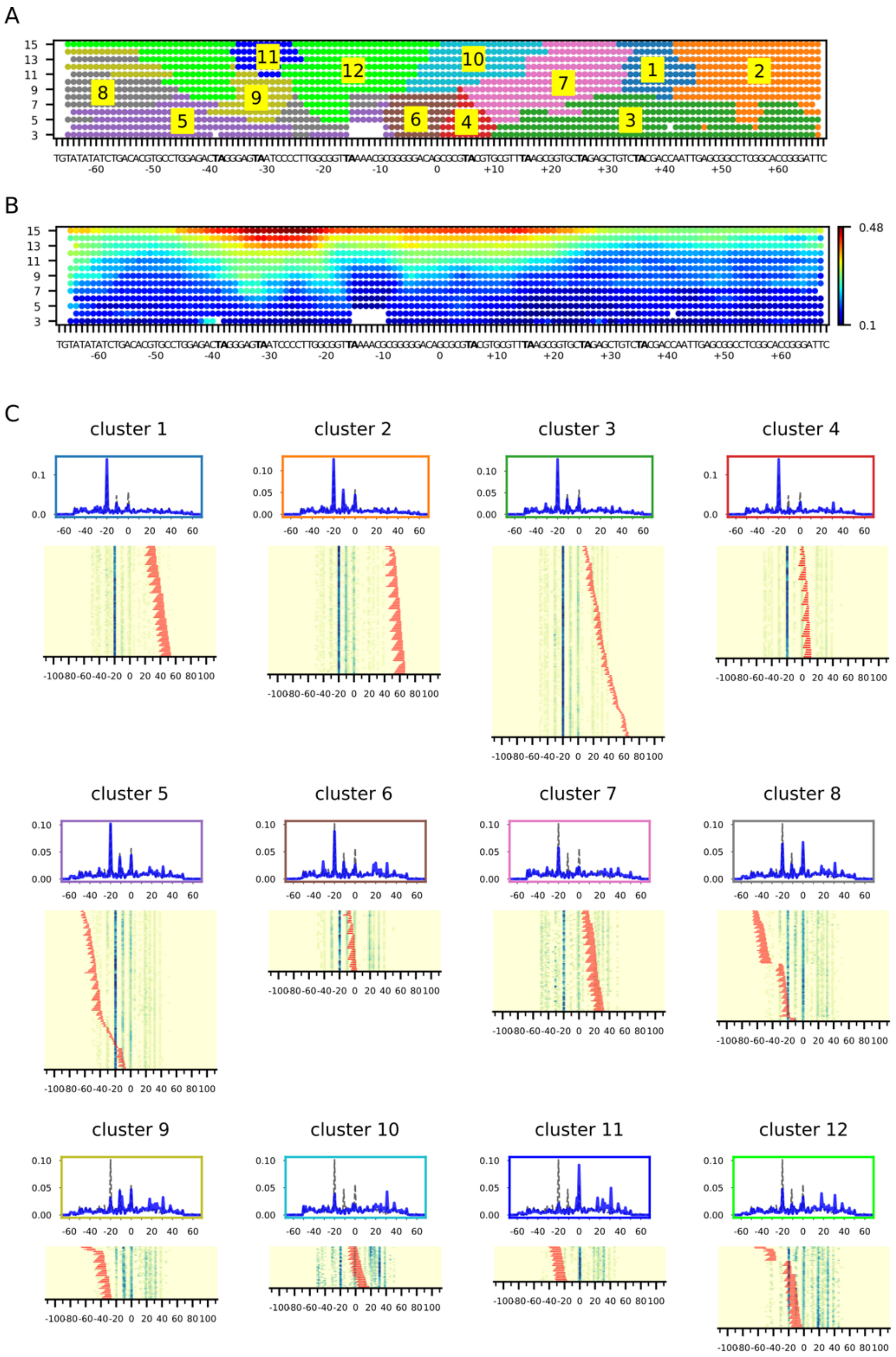


**Supplementary Figure S4. Clustering analysis of the poly(dA:dT) library based on nucleosome positions after sliding by Chd1.**

(A) The poly(dA:dT) library was clustered into 12 groups after sliding by Chd1. (B) A KL-divergence heatmap shows the positions and lengths of poly(dA:dT) tracts that most strongly altered the dyad pattern after Chd1 sliding. (C) Heatmaps of dyad positions grouped according to each cluster.

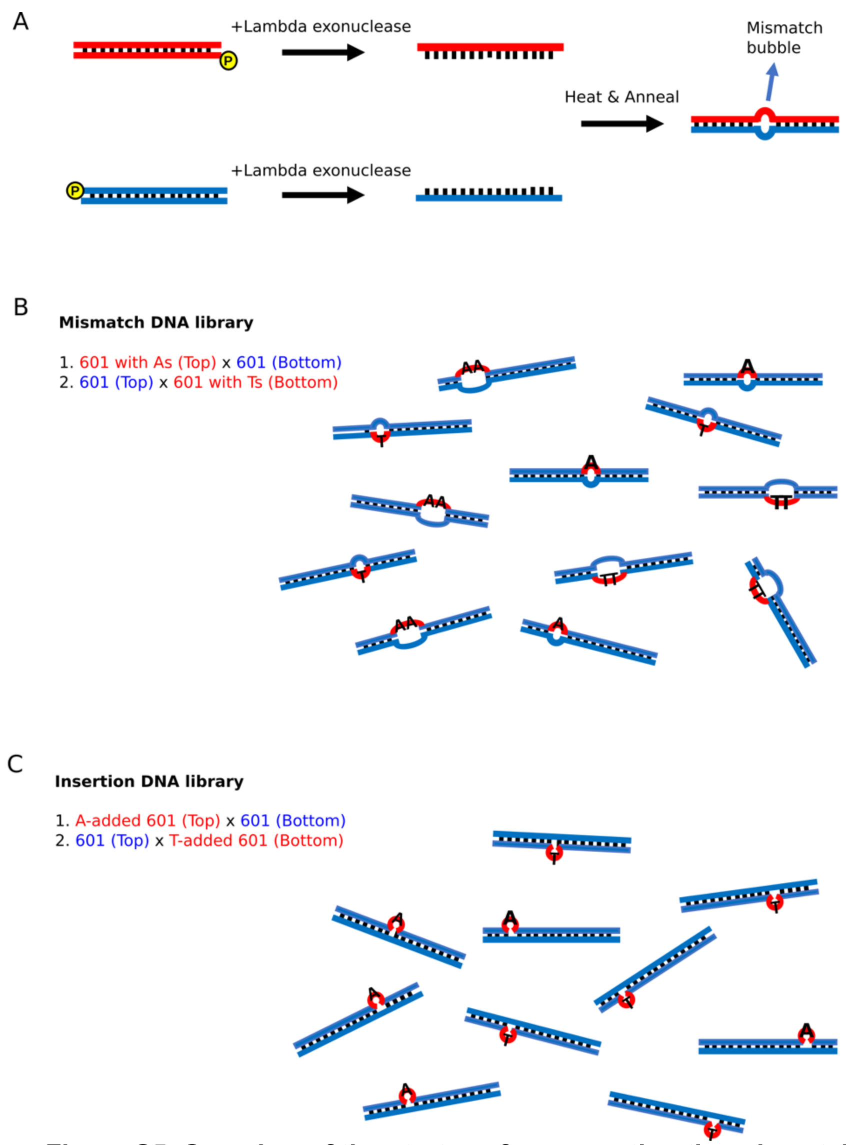


**Supplementary Figure S5. Overview of the strategy for generating the mismatch and single-insertion DNA library.**

(A) To generate mismatches or single-nucleotide insertions, the two strands of the duplex were produced in two separate reactions (red and blue). Lambda exonuclease, which has preferred activity toward strands with phosphorylated 5’ ends, was then added to selectively eliminate one strand. When the original templates differ at defined positions, annealing the resulting single strands produced duplexes containing mismatches or insertions. (B) The mismatch library is the mixture of two mismatch types: Adenine (A) substitutions in the top 601 strand paired with the original 601 bottom strand, and Thymine (T) substitutions in the bottom 601 strand paired with the original 601 top strand. (C) The insertion library is a mixture of two insertion types: Adenine (A) insertions in the top 601 strand paired with the original 601 bottom strand, and Thymine (T) insertions in the bottom 601 strand paired with the original 601 top strand.

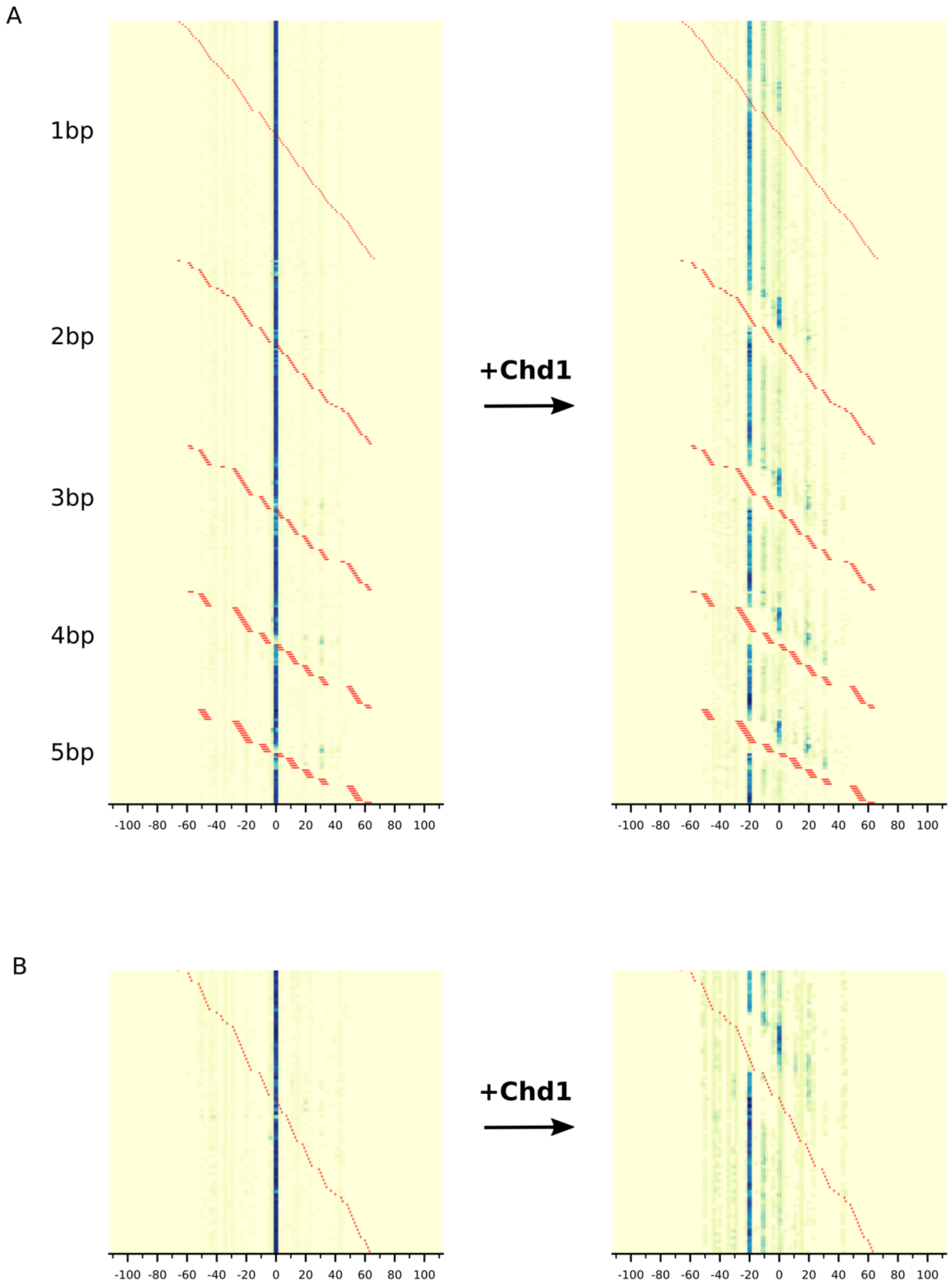


**Supplementary Figure S6. Heatmaps of dyad positions for mismatch and insertion library, before and after sliding by Chd1.**

(A) Heatmaps of mismatch library, grouped by mismatch position and length. (B) Heatmaps of single-insertion library, ordered by position. The location of mismatches and insertions are indicated by red bars.

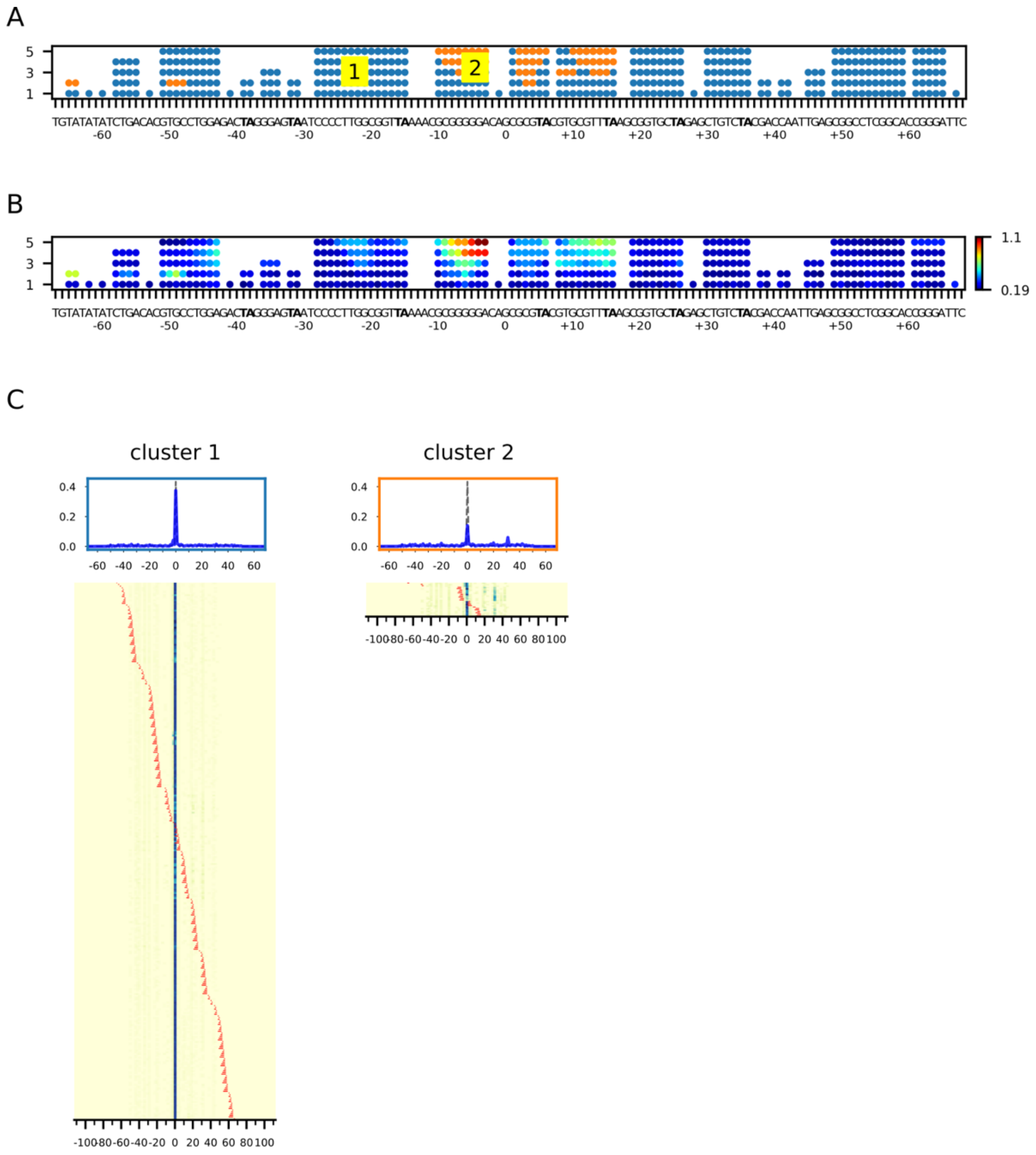


**Supplementary Figure S7. Clustering of mismatch library, before sliding by Chd1.**

(A) Based on the similarity of dyad positions and distributions, the mismatch library data was clustered into 2 groups. (B) A KL-divergence heatmap shows the most sensitive area in the Widom 601 DNA by perturbations. (C) Dyad positions for each mismatch are shown as heatmaps according to each cluster. Red bars indicate the length and position of mismatches.

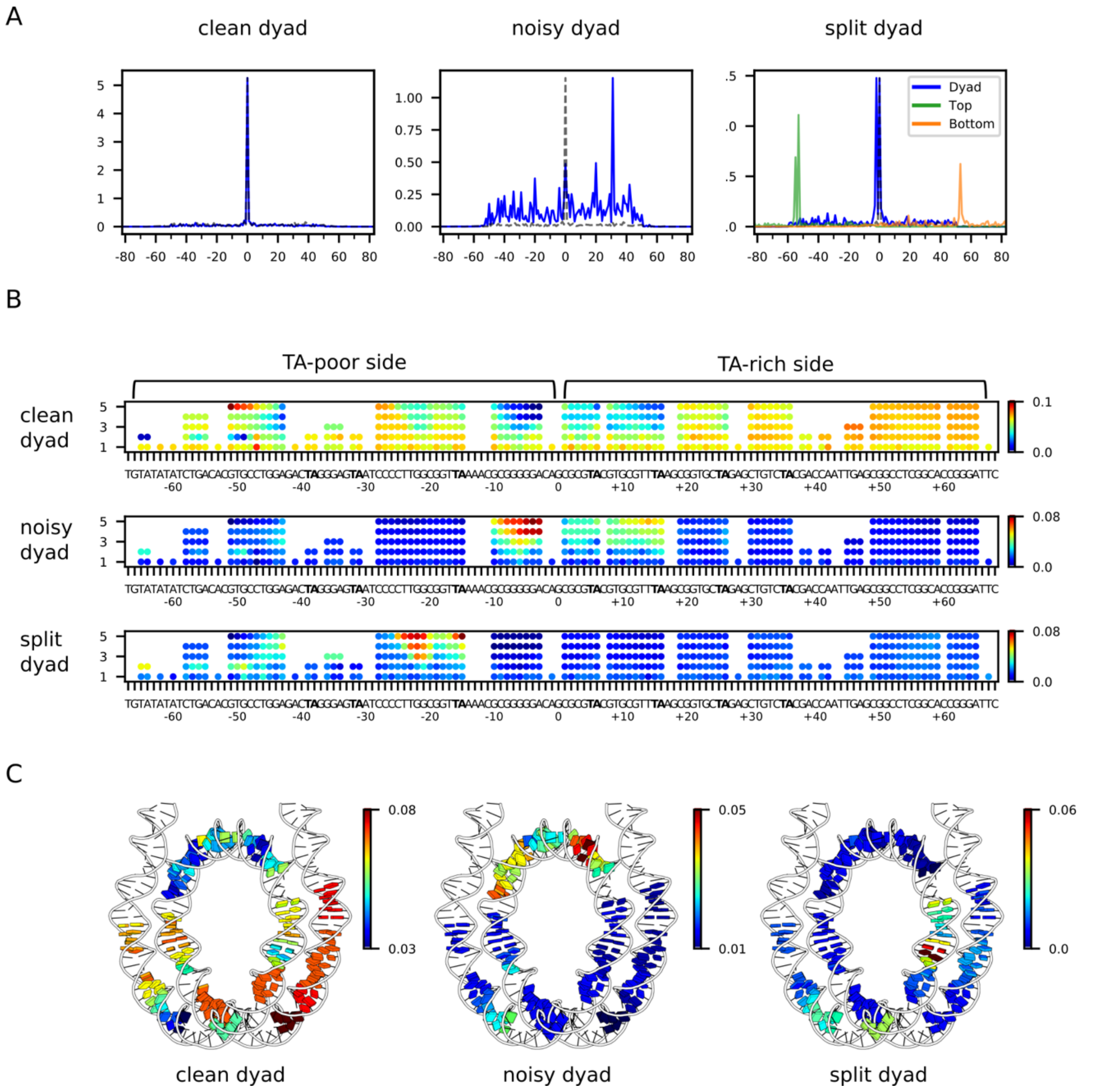


**Supplementary Figure S8. NMF analysis for nucleosome positioning on mismatch library before sliding by Chd1.**

(A) Through NMF analysis, all nucleosome positioning data was linearly decomposed into three basis patterns: clean dyad, noisy dyad, and split dyad. Calculated dyad positions are shown in blue, and sites of H2B(S53C) cross-linking are shown in orange and green. The canonical 601 dyad position is indicated by a dotted line. (B) For each basis pattern, the corresponding NMF scores were mapped into the Widom 601 sequence. (C) The NMF scores for 3 bp mismatches were mapped onto a nucleosome structure (6WZ5).

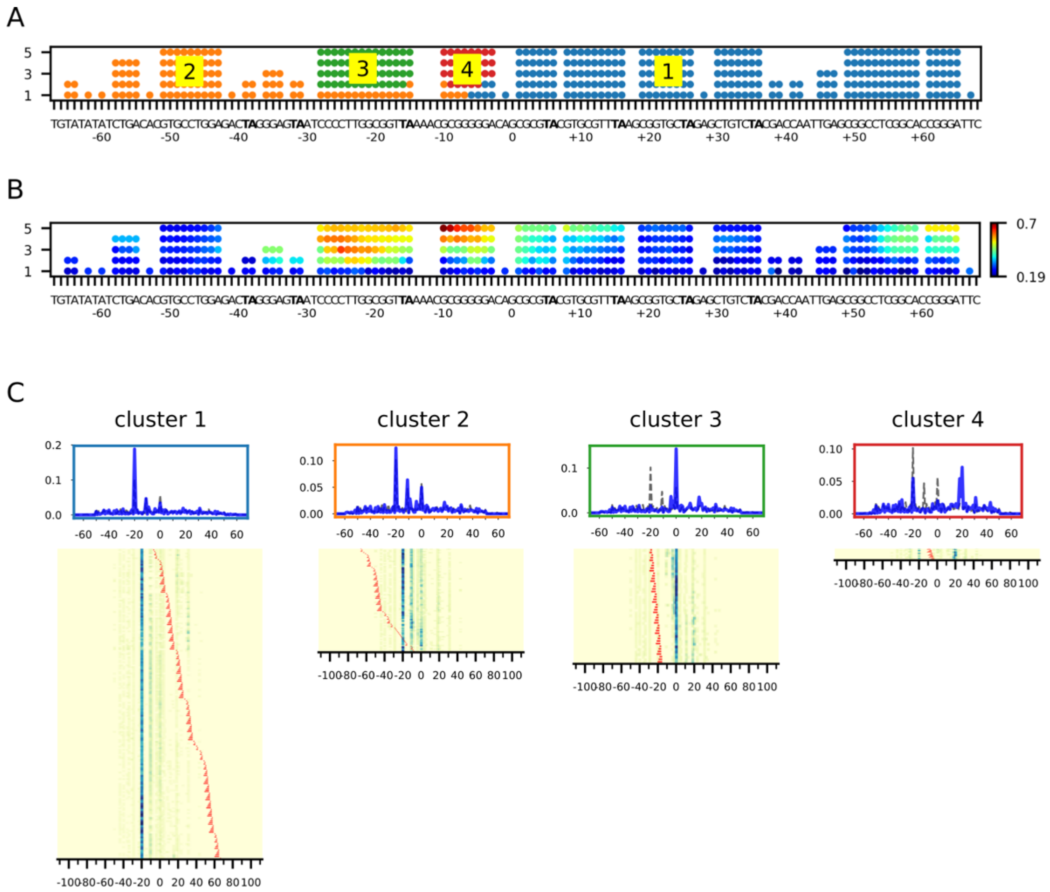


**Supplementary Figure S9. Clustering analysis of the 601 mismatch library after sliding by Chd1.**

(A) The mismatch library was clustered into 4 groups after sliding by Chd1. (B) A KL-divergence heatmap shows the positions and lengths of mismatches that most strongly altered the dyad pattern after Chd1 sliding. (C) Heatmaps of dyad positions grouped according to each cluster.

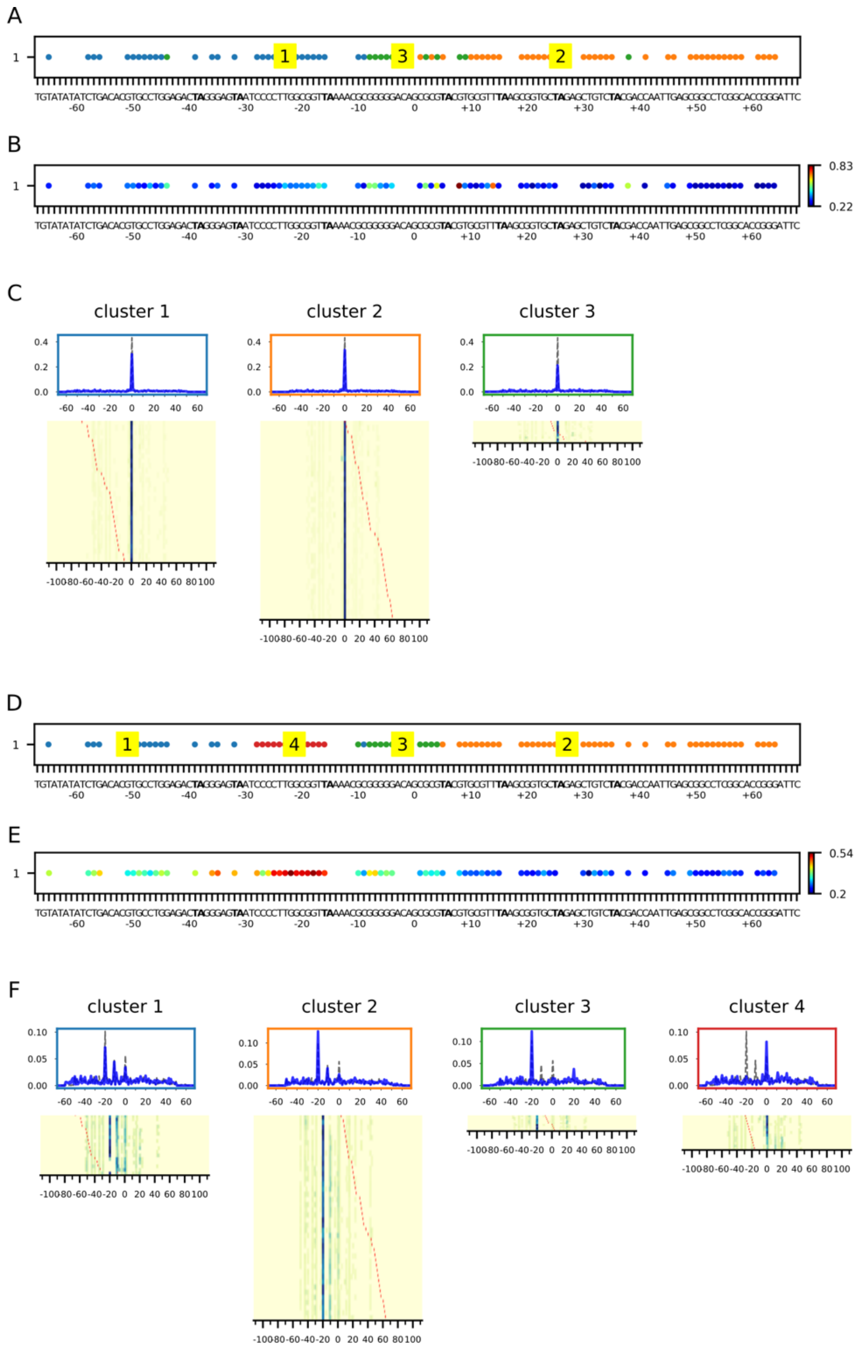


**Supplementary Figure S10. Clustering analysis of the 601 insertion library before and after sliding by Chd1.**

Through Spectral clustering analysis, the data were grouped and mapped on the Widom 601 before sliding (A) and after sliding (D) by Chd1. KL-divergence maps are show before sliding (B) and after sliding (E). For each cluster, the average positioning signal and data heatmap is shown for before sliding (C) and after sliding (F) by Chd1.

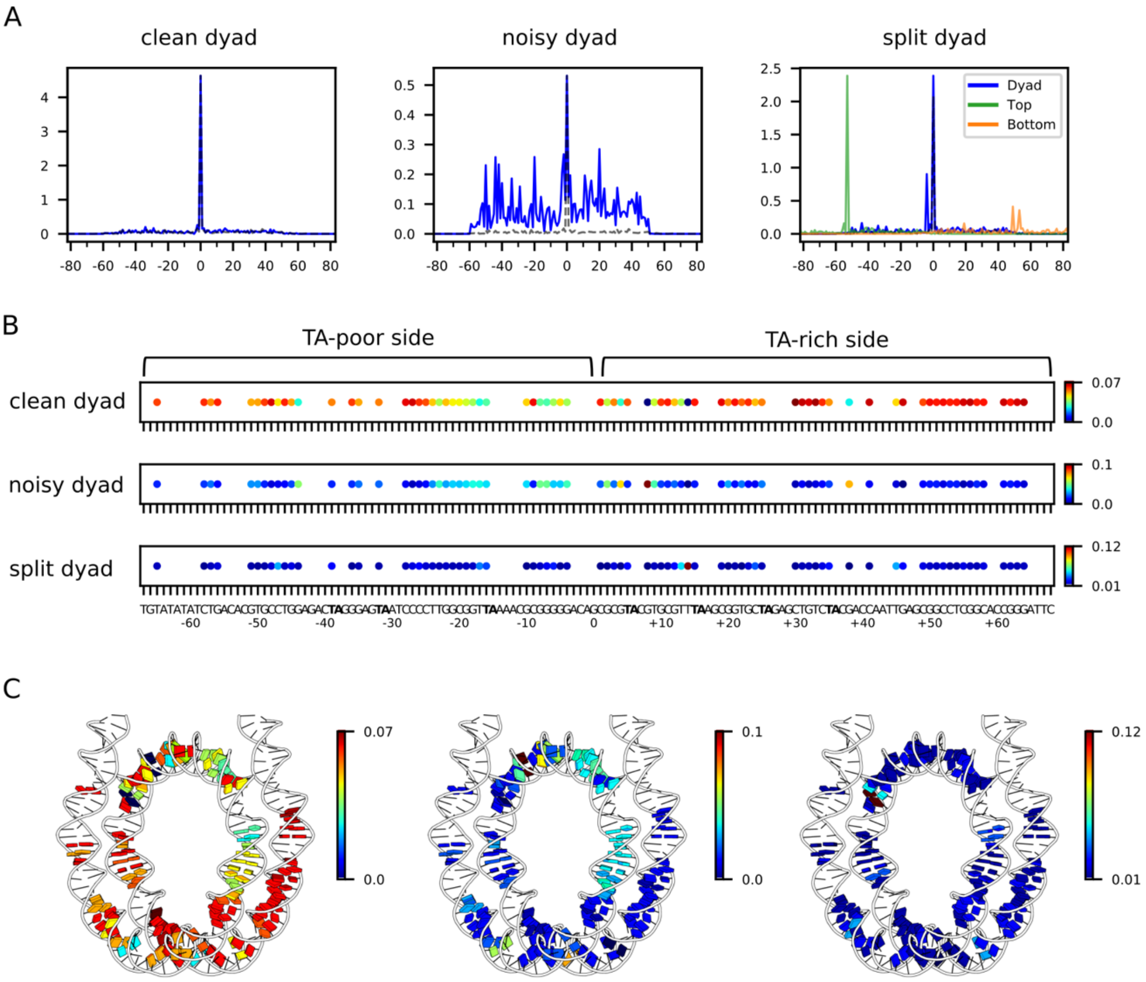


**Supplementary Figure S11. NMF analysis for nucleosome positioning on insertion library before sliding by Chd1.**

(A) Through NMF analysis, all nucleosome positioning data is linearly decomposed into three basis patterns: clean dyad, noisy dyad, and split dyad. Calculated dyad positions are shown in blue, and sites of H2B(S53C) cross-linking are shown in orange and green. The canonical 601 dyad position is indicated by a dotted line. (B) For each basis pattern, the corresponding NMF scores were mapped into the Widom 601 sequence. (C) The NMF scores were mapped onto a nucleosome structure (6WZ5).

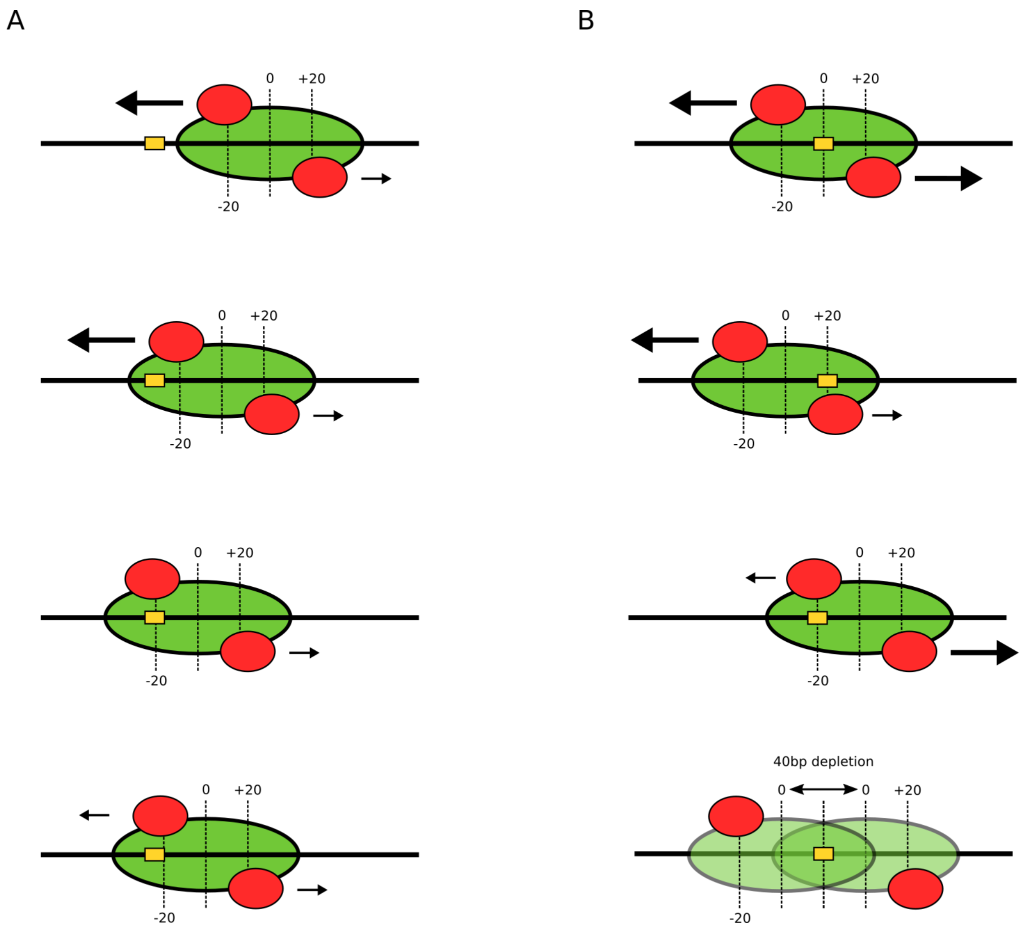


**Supplementary Figure S12. Models explaining nucleosome distributions through back-and-forth sliding by Chd1.**

Due to the two-fold symmetry of the nucleosome, there are two SHL2 sites where remodelers (red) can act. Each remodeler is positioned to move the nucleosome in opposite directions (arrows). (A) If a remodeler pulls a segment of DNA onto its own site that slows or stops sliding (yellow box), a remodeler acting at the other SHL2 can reverse the direction of DNA movement and shift that segment off of first site. (B) DNA segments that block remodelers can also cause depletion of certain nucleosome positions. When such segments (yellow box) are located between the two SHL2 sites, then either remodeler can initially move the site toward the other SHL2. Once reaching one SHL2 site, however, the DNA can no longer reverse, thus excluding dyad positions over a ~40 bp stretch.
